# CRISPR/Cas9-mediated Knock-in of an Optimized TetO Repeat for Live Cell Imaging of Endogenous Loci

**DOI:** 10.1101/162156

**Authors:** Ipek Tasan, Gabriela Sustackova, Liguo Zhang, Jiah Kim, Mayandi Sivaguru, Mohammad HamediRad, Yuchuan Wang, Justin Genova, Jian Ma, Andrew S. Belmont, Huimin Zhao

**Affiliations:** Department of Biochemistry, University of Illinois at Urbana-Champaign, Urbana, IL 61801, USA; Department of Cell and Developmental Biology, University of Illinois at Urbana-Champaign, Urbana, IL 61801, USA; Carl R. Woese Institute for Genomic Biology, University of Illinois at Urbana-Champaign, Urbana, IL 61801, USA; Department of Chemical and Biomolecular Engineering, University of Illinois at Urbana-Champaign, Urbana, IL 61801, USA; Computational Biology Department, School of Computer Science, Carnegie Mellon University, Pittsburgh, PA 15213, USA; Center for Biophysics and Quantitative Biology, University of Illinois at Urbana-Champaign, Urbana, IL 61801, USA; Department of Chemistry, University of Illinois at Urbana- Champaign, Urbana, IL 61801, USA; Department of Bioengineering, University of Illinois at Urbana-Champaign, Urbana, IL 61801, USA

## Abstract

Nuclear organization has an important role in determining genome function; however, it is not clear how spatiotemporal organization of the genome relates to functionality. To elucidate this relationship, a high-throughput method for tracking any locus of interest is desirable. Here, we report an efficient and scalable method named SHACKTeR (Short Homology and CRISPR/Cas9-mediated Knock-in of a TetO Repeat) for live cell imaging of specific chromosomal regions. Compared to alternatives, our method does not require a nearby repetitive sequence and it requires only two modifications to the genome: CRISPR/Cas9-mediated knock-in of an optimized TetO repeat and its visualization by TetR-EGFP expression. Our simplified knock-in protocol, utilizing short homology arms integrated by PCR, was successful at labeling 9 different loci in HCT116 cells with up to 20% efficiency. These loci included both nuclear speckle-associated, euchromatin regions and nuclear lamina-associated, heterochromatin regions. We anticipate the general applicability and scalability of our method will enhance causative analyses between gene function and compartmentalization in a high-throughput manner.

## Introduction

It is becoming increasingly clear that spatiotemporal organization of the mammalian genome within the nucleus is not random, but is highly regulated (Dixon et al., 2012; Misteli, 2007). Although nuclear architecture is suggested as an important regulator for various nuclear processes including gene expression, the link between subnuclear localization and gene function remains elusive. To relate genome function to higher order nuclear organization, a direct, microscopy-based method for live cell tracking of the dynamics of any specific endogenous locus of interest is necessary.

DNA fluorescence *in situ* hybridization (DNA-FISH) is a commonly used method for imaging a specific region within the chromosome but is technically challenging. Because of the harsh DNA denaturation conditions required, structural preservation is poor, yet increasing fixation strength to counteract this structural perturbation results in decreased detection efficiency. DNA-FISH also frequently has high background with both false positive and false negative rates. For all of these reasons, DNA-FISH is a difficult, low efficiency visualization method, which also has serious intrinsic resolution limitations due to problems with structural preservation of chromosome structure. Finally, DNA-FISH is performed in fixed cells, and therefore incompatible with tracking dynamics of DNA.

Live cell imaging of DNA in mammalian cells was previously carried out by using a fluorescent repressor-operator system (FROS) (Belmont, 2001; Robinett et al., 1996), using enrichment of a fluorescent protein (FP) at a specific site on the DNA to produce a fluorescent signal that is above the background noise. For this purpose, FPs such as green fluorescent protein (GFP) were fused to DNA binding proteins that recognized a repeating DNA sequence. In the two most commonly used systems, repeating sequences of Lac operators (LacO) or Tet operators (TetO) are used as a DNA tag and FP-fused Lac repressor (LacI) or Tet repressor (TetR), respectively, are used for visualization of the tag. In most previous examples, live cell imaging has focused on plasmid or BAC integrated transgenes, although the behavior of these transgenes may not fully recapitulate the behavior of the endogenous locus. In a more recent example, these operator arrays have been targeted to endogenous chromosome loci using homologous recombination (HR) (Masui et al., 2011). However, in this previous work a low targeting efficiency was observed, indicating the need for new targeting strategies with higher efficiency.

Discovery of novel modular proteins whose DNA recognition specificity can be easily tailored lead to the development of alternative approaches for visualizing endogenous loci. Transcription activator-like effectors (TALEs) contain repeats of a 33–35 amino acid sequence, which only differ at the 12th and 13th amino acids, determining the base specificity of each repeat (Boch et al., 2009; Moscou and Bogdanove, 2009). Clustered regularly interspaced short palindromic repeats (CRISPR)/CRISPR-associated protein 9 (Cas9) includes an sgRNA that can recognize a 20 nt DNA sequence based on Watson-Crick base pairing and the sgRNA recruits the Cas9 endonuclease, associated with the scaffold of sgRNA, to the target DNA sequence (Ran et al., 2013). If the target sequence is followed by a protospacer adjacent motif (PAM), which is 5’-NGG-3’ for Cas9 from *Streptococcus pyogenes*, then Cas9 can induce a double strand break (DSB) approximately 3 nucleotides upstream of the PAM. Coupled with fluorescent tags, both TALEs and a catalytically inactive version of Cas9 (dCas9) were used for visualization of naturally occurring repetitive sequences within mammalian genomes (Chen et al., 2013; Ma et al., 2013; Miyanari et al., 2013; Thanisch et al., 2014). However, repetitive sequences are not equally distributed throughout the genome, necessitating other strategies for visualizing non-repetitive regions. Visualization of non-repetitive sites by TALEs or dCas9 requires introduction of a set of many TALEs or sgRNAs recognizing multiple sites within the target region, which could be technically challenging and inefficient.

In addition to its above-mentioned use for tracking of genes in mammalian cells, increasing the efficiency of homology-directed genome editing is among one of the most commonly used applications of the CRISPR/Cas9 system. In mammalian cells, non-homologous end joining (NHEJ) is the predominant mechanism for repairing DSBs created by CRISPR/Cas9. NHEJ is an error-prone mechanism that can cause insertion/deletion mutations; hence, it is mostly used for gene knockout. Alternatively, in the presence of a DNA that is homologous to the sequences flanking the DSB site, such as an exogenous donor DNA with suitable homology arms, the DSB can be repaired by HR as well (Johnson and Jasin, 2001). Depending on the design of the exogenously delivered donor vector, gene knock-in, deletion or replacement can be achieved via homology-directed repair of the DSBs. CRISPR-mediated knock-in has been used for various applications including epitope or GFP tagging of endogenous proteins (Park et al., 2014; Ratz et al., 2015; Savic et al., 2015). One application of CRISPR-mediated knock-in that has not been explored yet is the tagging of the DNA itself for visualizing the position and dynamics of specific chromosomal regions.

Here, we developed an easy, efficient and scalable method named SHACKTeR (short homology arm and CRISPR/Cas9 mediated knock-in of a TetO repeat) for tagging and live cell imaging of non-repetitive, endogenous chromosome regions. With the help of an improved donor DNA design and an optimized, irregular TetO repeat, it was possible to achieve targeted insertion efficiencies of up to 20%. We demonstrated the general applicability of SHACKTeR by inserting the TetO repeat into 9 different chromosome regions, representing both gene-rich, transcriptionally active and nuclear speckle-associated sites as well as gene-poor, transcriptionally inactive and nuclear lamina-associated sites. TetO repeats as short as 48 were successfully visualized with high efficiency using lentiviral TetR-EGFP expression. Our labeling strategy was highly specific as demonstrated by the lack of additional off-target spots in the nucleus. The ease-of-use and the scalability of SHACKTeR will allow us to label large numbers of endogenous chromosome loci for both fixed-cell and live-cell imaging, which we expect will lead to an improved understanding of the relationship between gene compartmentalization and function.

## Results

### Optimized TetO multimers enable PCR-based creation of a linear donor DNA with short homology arms

A major bottleneck in the traditional HR method is the use of long homology arms of several kilobases, which need to be PCR-amplified from the genomic DNA (gDNA) and then cloned into a donor vector. Constructing these donor vectors impedes high throughput applications exploiting HR-mediated targeting. Based on previous work using limited homology arms as short as 50 base pairs (Orlando et al., 2010), we devised a one-step, PCR-based method to create a linear donor DNA containing a TetO repeat. We amplified the TetO repeat and selection marker by PCR using primers which included 50 nt 5’ extension sequences homologous to the target site sequence (Figure 1A). The PCR product could then be gel or PCR-purified and used directly for transfection as a donor construct with 50 bp homology arms. For creating all future donor constructs we first created a plasmid to use as a PCR template.

**Figure 1:**
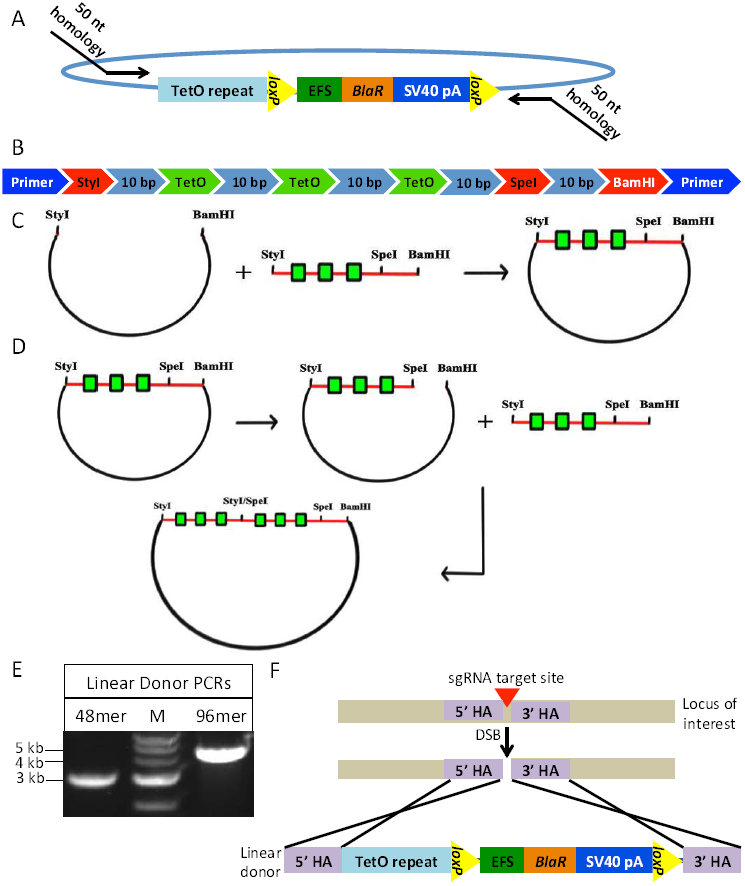
CRISPR/Cas9-mediated knock-in of an optimized TetO repeat using linear donor DNA with short homology arms. **(A)** Schematic of the strategy to create linear donor DNA with short homology arms via PCR. The final plasmid used as a template is shown together with the primers used for amplification of the linear donor DNA (black arrows). Blasticidin resistance (*BlaR*) gene under EFS promoter was used for positive selection. The selection cassette is flanked with *loxP* sites in case it needs to be removed. **(B)** Schematic of the complete initial cassette containing three TetO repeats (green), four random 10 bp (light blue) sequences, three restriction sites StyI, SpeI, and BamHI (all of them are in red), and primers (dark blue) used for synthesis of the complementary strand and amplification of the product. **(C)** Schematic of the insertion of 3mer TetO cassette into pSP2 vector through StyI and BamHI sites. Each TetO sequence is indicated with a green box. **(D)** Schematic of the assembly of the TetO repeat multimers. StyI/SpeI site, created by insertion of the TetO cassette, cannot be cut by either of these two enzymes. **(E)** Results of the PCRs to amplify 48-mer and 96-mer TetO repeat donors for the *HSP70* locus. The expected band sizes were around 3 kb for the 48-mer and around 4.6 kb for the 96-mer TetO donor. NEB 1 kb ladder (M) was used. **(F)** Schematic of the knock-in strategy. Homology arms (HA) are shown in purple. sgRNA target site is indicated with a red triangle. DSB: double strand break.

In previous studies, perfect direct repeats of TetO or LacO sequences were used as tagging sequences. Such direct repeats, however, create significant plasmid instability when propagated in bacteria, making all downstream applications difficult and time-consuming, as multiple clones must be grown and screened to verify that the direct repeats remain intact. Moreover, PCR-amplification of direct repeats is challenging (Hommelsheim et al., 2014).

To overcome these problems associated with direct operator repeats, we created irregular 48-mer and 96-mer TetO repeats. These repeats contained the 19 bp TetO sequence separated by random 10 nt sequences. Since RecA pairing in bacteria requires at least 25 nt of homology, the presence of 10 nt random sequences between TetO repeats reduces recombinational and replicational instability of these sequences within plasmids grown in bacteria. A similar approach was used to construct irregular Lac and Tet operator repeats which had increased stability when propagated in bacteria (Lau et al., 2003). We anticipated that this irregular TetO repeat would also facilitate PCR-amplification of this repetitive sequence.

We also purposely designed these 48-mer and 96-mer TetO repeats to contain no CpG dinucleotides that could be templates for DNA methylation after introduction into mammalian cells. We rationalized that the absence of DNA methylation would reduce the tendency of these TetO arays to form heterochromatin after integration into the mammalian chromosome. The Tet operator itself lacks CpG dinucleotides. We used a mixture of only three nucleotides-G, T, and A-to introduce random 10 nt sequences between the TetO sequences. A 50:25:25 ratio of G, T, and A nucleotides was used to produce a 50% GC content for these 10 nt spacer sequences. The cloning strategy we used was designed to produce TetO arrays in which all 10 bp spacer sequences were unique; this was confirmed by sequencing the final 48-mer and 96-mer constructs.

Although 96-mer TetO repeats were used previously for visualization (Normanno et al., 2015), we predicted the shorter 48-mer TetO array might produce higher knock-in efficiency. Moreover, we reasoned that insertion of a 48-mer repeat into the genome might be less perturbing compared to introduction of a 96-mer or longer repeat. The plasmids containing these 48-mer and 96-mer TetO repeats were further modified by cloning a blasticidin resistance cassette flanked by *loxP* sites (Figure 1A). The overall cloning strategy we used to create these TetO arrays is illustrated in Figure 1 B,C,D. As anticipated, the optimized, irregular nature of these TetO repeats enabled easy PCR amplification of both 48-mer and 96-mer TetO donors as verified by visualization of a single, clear PCR-product band after gel electrophoresis (Figure 1E).

### Comparison of the knock-in efficiencies of 48-mer and 96-mer TetO repeats

To compare the efficiencies of 48-mer versus 96-mer TetO knock-ins, we targeted the heat shock protein 70 (*HSP70*) human chromosome locus containing three HSP70 homologous genes (*HSPA1A, HSPA1B, HSPA1L*). We used the genetically stable, near diploid human HCT116 colorectal carcinoma cell line (Haigis et al., 2002) for all of our targeted insertions. Using a near diploid cell line made genotyping results and the specificity of our targeting easier to interpret.

Since homology-directed repair of DSBs is normally inefficient in mammalian cells (Mao et al., 2008a, 2008b), our SHACKTeR strategy uses the CRISPR/Cas9 system to increase the efficiency of the TetO repeat HR knock-in (Figure 1F). We designed an sgRNA targeting an intergenic region approximately 16 kb downstream of the *HSPA1B* gene. Enhancer-like or promoter-like regions as predicted from the ENCODE data in HCT116 cells were avoided for targeting by CRISPR/Cas9. Transcription factor binding sites were also avoided. An sgRNA target site with a unique and non-repetitive sequence, based on the RepeatMasker track in UCSC Browser, was chosen for better specificity. We also avoided inclusion of repetitive or low complexity sequences within the 50 nt homology arms to minimize off-target integration.

Linear 48-mer and 96-mer TetO donors were PCR-amplified and gel-purified (Figure 1E), followed by co-transfecting each donor into HCT116 cells together with the CRISPR/Cas9 plasmid at a 1:1 molar ratio. This CRISPR/Cas9 plasmid contained expression cassettes for both the sgRNA targeting the *HSP70* locus and the Cas9 protein. As a negative control for CRISPR/Cas9-mediated HR knock-in, we also transfected the cells with 48-mer or 96-mer TetO donor DNA only. After one week of positive selection with blasticidin, mixed pool cells were collected to determine the presence of cells with successful knock-in using the specific PCR-amplification of the knock-in allele as the assay (Figure 2A). We detected successful integration into the target site of both 48-mer and 96-mer TetO inserts and these integrations were dependent on the co-transfection of the CRISPR/Cas9 plasmid (Figure 2A). The number of blasticidin-resistant cells observed after transfection in the absence of the CRISPR/Cas9 plasmid, due to random integration of the donor through NHEJ, was noticeably lower than that observed with co-transfection with the CRISPR/Cas9 plasmid (data not shown).

**Figure 2:**
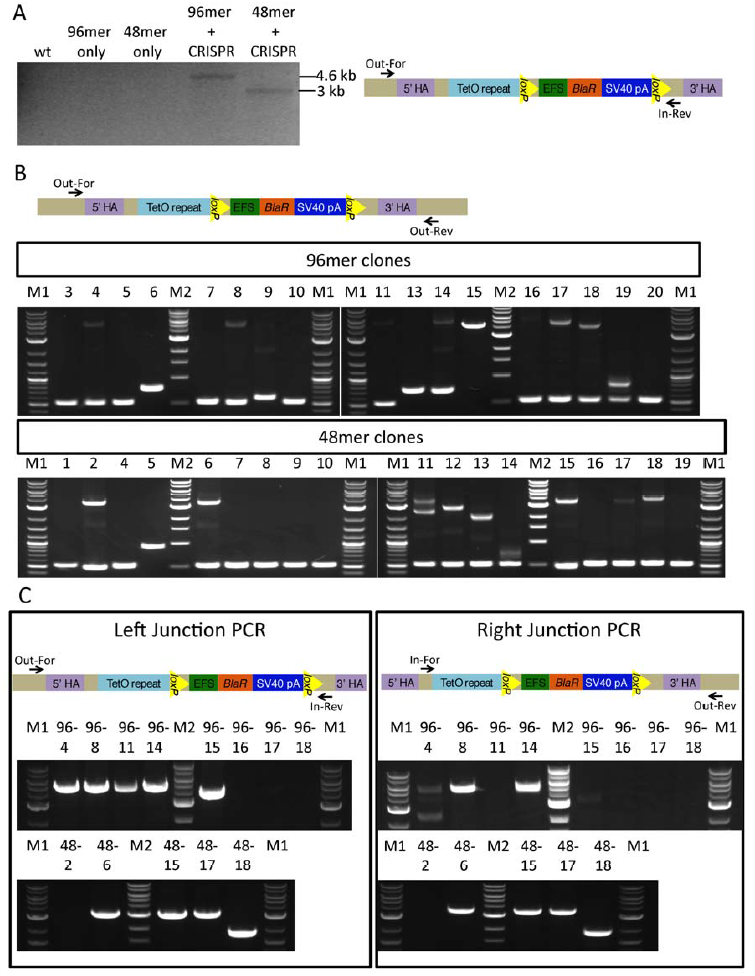
Comparison of 48-mer and 96-mer TetO knock-in. **(A)** PCR analysis of mixed pool cells that were collected after transfection of the CRISPR/Cas9 plasmid together with either 48-mer or 96-mer TetO donor DNA. Cells were collected for genotyping 7 days after blasticidin selection. Results from transfection with 48-mer or 96-mer TetO donor-only were used for controlling CRISPR/Cas9-mediated knock-in. Schematic of the primers used for genotyping is shown on the right. Out-For and In-Rev primers were used for the PCR-amplification of the knock-in band only. The band was expected to be around 3 kb for 48-mer TetO knock-in and 4.6 kb for 96-mer TetO knock-in. PCR from the gDNA of wild-type (wt) HCT116 cells was used as a negative control. **(B)** Genotyping of clones from 48-mer or 96-mer TetO knock-in. Out-For and Out-Rev primers (shown on top) were used for the genotyping PCR. These primers anneal to the gDNA sequence outside of the homology arms. Expected band for the clones with the correct knock-in is around 3.4 kb for 48-mer TetO and around 5 kb for 96-mer TetO knock-in. Clones 96-1, 96-2, 96-12, and 48-3 were not included in genotyping due to their growth deficiency. **(C)** Left and right junction PCR results for the 96-mer (top gels) and 48-mer (bottom gels) knock-in clones identified as potentially correct from panel B. Out-For and In-Rev primers were used for left junction PCR, and In-For and Out-Rev primers were used for right junction PCR. Locations of the primers for each PCR is shown on top. Expected left junction PCR band sizes for the correctly targeted clones are around 3 kb for 48-mer TetO and around 4.6 kb for 96-mer TetO knock-in. Expected right junction PCR band sizes for the correctly targeted clones are around 3.4 kb for 48-mer TetO and around 5 kb for 96-mer TetO knock-in. NEB 2-log (M1) and 1kb ladder (M2) were used.

To determine knock-in efficiency, single clones were isolated by serial dilution. We excluded all clones that showed noticeably slower rates of proliferation. In total, we analyzed 17 clones from the TetO 96-mer knock-in and 18 clones from the 48-mer knock-in. Genotyping of isolated clones was first performed using a pair of primers that both annealed to the gDNA outside of the homology arms (Figure 2B). As a result, both knock-in and non-knock-in alleles could be amplified, showing whether both, none or only one of the alleles were modified.

As a secondary assay, we performed PCRs to amplify 5’ and 3’ junctions of the knock-in cassette (Figure 2C) in the subset of single cell clones that had shown insertions at the targeted site. In these junction PCRs, one primer anneals to gDNA outside of the homology arm and the other one anneals to the insert. This secondary assay determines whether the donor was inserted in correct direction. As a final test for the clones that gave correct results in the three previously described genotyping PCRs, we gel-purified the left junction PCR band and digested it with SpeI restriction enzyme, which cuts between the TetO repeat and the selection marker (Figure 2-figure supplement 1A). This digestion is helpful in assaying the integrity of TetO repeat. Based on this genotyping, 3/18 of the 48-mer clones (48-6, 48-15 and 48-17) and 2/17 of the 96-mer clones (clones 96-8 and 96-14) showed correct targeted insertions. All of these clones showed heterozygous TetO insertions (Figure 2B).

An unexpected result of this genotyping was the observation of PCR bands whose length was longer than the band expected from the unmodified wild-type (wt) allele but shorter than the band expected from a full-length knock-in allele. We suspected that the TetO repeats might be subject to shortening during HR. This shortening was confirmed by sequencing and SpeI digestion-test of the left junction PCR bands from clones 48-11 and 48-12 which showed shortened TetO repeats but an intact selection cassette (Figure 2-figure supplement 1B). Similar shortening of lacO repeats has been observed during P element transposition in *Drosophila* (A. Belmont, unpublished data). Because donor DNA was size selected by gel-purification, it is unlikely that these shortened TetO arrays arose during bacterial growth of the donor template plasmid or PCR. Because a full-length TetO array integrated correctly at the target site was stable in length after more than a month of continuous HCT116 cell passaging (data not shown), it is therefore also unlikely that this shortening of the TetO arrays occurred after integration. Therefore, we infer this shortening results from recombination of repetitive sequences during the HR process, suggesting our knock-in strategy might have shown even higher efficiency with a non-repetitive insert.

To test whether 48 TetO repeats were sufficient for visualization of the *HSP70* locus, clone 48-15 was transfected with the p3’SS TetR-EGFP plasmid expressing the Tet repressor fused to EGFP. Two days after transfection, mixed pool cells were analyzed using fluorescence microscopy. In most of the cells expressing TetR-EGFP, we observed a single, clear EGFP-tagged spot in the nucleus (Figure 2–figure supplement 1C), consistent with the heterozygous insertion demonstrated by genotyping. This microscopy observation of a single spot goes beyond the genotyping results in showing the absence of additional off-target insertions. In some cells there were two nearby spots, possibly due to replication of the locus. We could not see a spot in some of the cells due to either too high or too low TetR-EGFP expression.

### Scalability of the 48-mer TetO knock-in: targeting 9 additional chromosome loci with varying chromatin marks and sub-nuclear localizations

In subsequent experiments we used the 48-mer TetO, due to the possibility of a higher targeting efficiency with the shorter insert and, more importantly, the expectation that the smaller repeat size would be less likely to perturb the normal chromatin structure at the target insertion sites.

Recent work has suggested that the cutting efficiency of CRISPR/Cas9 may depend on the chromatin structure of the target site, with less efficient cutting observed at heterochromatic sites (Chen et al., 2016). Moreover, previous work showed that HR efficiency also can be affected by the transcriptional and chromatin status of the region, with active gene regions with euchromatic histone marks showing higher HR efficiency (Aymard et al., 2014). Thus, in addition to the *HSP70* locus that is known to be highly active in most of the cancer cells (Murphy, 2013), we wanted to further test the efficiency of our knock-in method targeting both euchromatic and heterochromatic target loci localized in different nuclear compartments.

We used a new method, TSA-Seq (Chen, 2016), to guide selection of target chromosome loci showing different nuclear compartment localization. Using TSA-Seq, the relative cytological distances of chromosomes genome-wide relative to the nuclear lamina and nuclear speckles was estimated in human K562 cells. Based on this TSA-mapping, we selected 4 chromosome regions (1, 2, 4 and 5) that were expected to be nuclear speckle-associated (data not shown). Nuclear speckles are enriched in pre-mRNA splicing factors and mostly interact with active gene regions (Hall et al., 2006; Shopland et al., 2003; Spector and Lamond, 2011). For instance, *HSP70* locus (region 3) and HSP70 transgenes were known to interact with nuclear speckles when active (Jolly et al., 1999; Khanna et al., 2014). In K562 cells, the *HSP70* locus also was predicted to be speckle-associated by TSA-Seq analysis. As expected, these 5 nuclear speckle-associated sites all corresponded to gene-rich and transcriptionally active regions in both K562 and HCT116 cells based on ENCODE RNA-Seq data (Figure 3-figure supplement 1).

**Figure 3:**
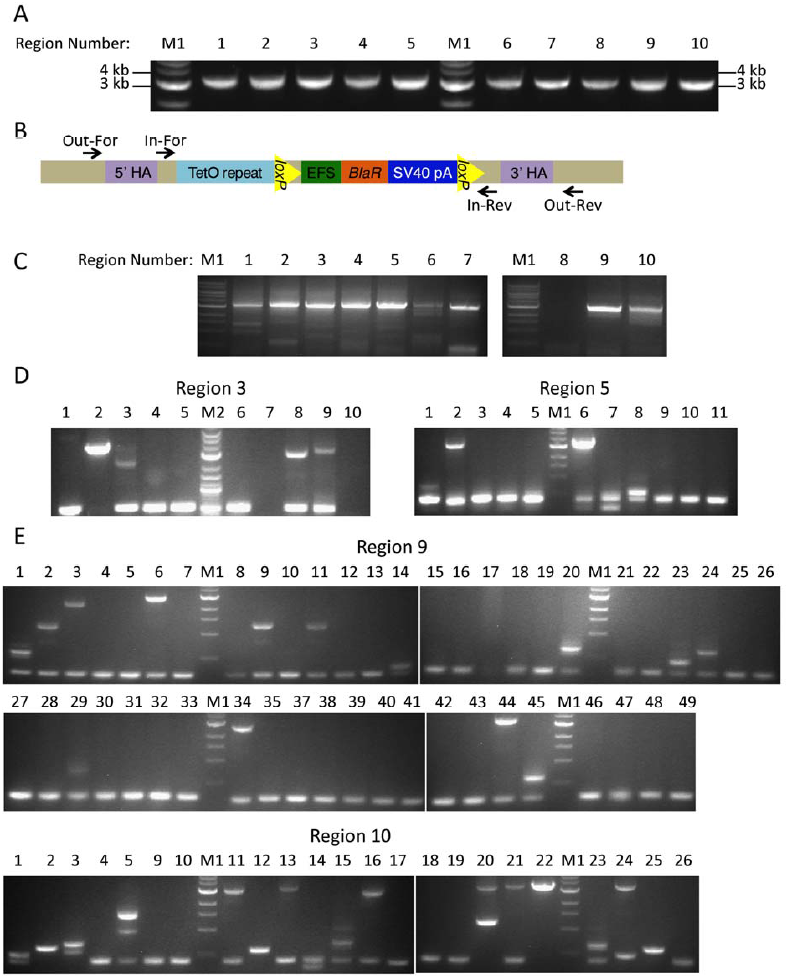
48-mer TetO knock-in into various loci and genotyping results. **(A)** Results of the PCRs for creating linear 48-mer TetO donor DNA for the knock-in into 10 different regions. The donors are expected to be around 3.1 kb. **(B)** Schematic of the primers used for genotyping. **(C)** PCR analysis of mixed pool cells after transfection of the CRISPR/Cas9 plasmid and 48-mer TetO donor DNAs for knock-in into 10 different regions. Out-For and In-Rev primers, described in panel B, were used for the PCR-amplification of the knock-in band only. Knock-in band is expected to be around 3.1 kb for regions 3, 8 and 9; around 3.2 kb for regions 4, 5, 7 and 10; around 3.3 kb for region 2 and 6; and 3.6 kb for region 1. **(D)** Genotyping of the clones from 48-mer TetO knock-in into regions 3 and 5. Out-For and Out-Rev primers were used for the genotyping PCR. Expected band sizes from the knock-in alleles are around 3.5 kb for region 3 and 3.4 kb for region 5. Expected bands from the wt alleles are 528 bp for region 3 and 389 bp for region 5. **(E)** Genotyping of the clones from 48-mer TetO knock-in into regions 9 and 10. Out-For and Out-Rev primers were used for the genotyping PCR. Expected band sizes from the knock-in alleles are around 3.3 kb for region 9 and 3.4 kb for region 10. Expected bands from the wt alleles are 290 bp for region 9 and 407 bp for region 10. M1: NEB 1 kb ladder. M2: NEB 2-log ladder.

Alternatively, we selected 4 chromosome regions (6-9) that were predicted to associate with the nuclear lamina in K562 cells based on both lamin TSA-Seq and lamin B1 DamID data (data not shown). Lamina-associated sites correspond to gene-poor, mostly inactive chromatin domains (Guelen et al., 2008; Kind and van Steensel, 2010; van Steensel and Belmont, 2017). These 4 regions all showed low levels of transcriptional activity in both K562 and HCT116 cells based on ENCODE RNA-Seq data (Figure 3-figure supplement 1). Additionally, we selected the β -globin locus (*HBB*) as our 10^th^ targeting region. Although in K562 cells this locus is active and not associated with the lamina, in other tissues and cell lines in which this locus is inactive, the β-globin locus associates with the nuclear periphery, or centromeric heterochromatin in some cell types (Hepperger et al., 2008; Ragoczy et al., 2006). Surprisingly, region 10 has high transcriptional activity in HCT116 cells (Figure 3-figure supplement 1), but is still associated with the lamina as determined by DNA-FISH. BAC (bacterial artificial chromosome) clones corresponding to all 10 regions and their coordinates can be found in Figure 3-figure supplement 2.

To knock-in an insert for the purpose of labeling a locus, the high compaction of mammalian chromatin introduces a considerable flexibility in terms of choosing an appropriate target sequence. Using FISH, hybridization of BAC probes of ~ 200 kb still appear as a near diffraction-limited spot by wide-field light microscopy; therefore a distance of up to ~ 100 kb from the target locus will locate at the same location as resolved by conventional fluorescence microscope. To find the best sgRNA for targeting each of the 9 additional regions, first we chose DNA regions which did not contain any exons to avoid disrupting coding sequences. The complete criteria used to choose sgRNAs is detailed in the Materials & Methods section. Briefly, in addition to the standard criteria for sgRNA design, we also avoided targeting regions with a repetitive or low complexity sequence within 50 nt upstream or downstream of the expected cut site, allowing us to design homology arms with high specificity. Potential sgRNAs with high scores were further evaluated to exclude any that targeted promoter or enhancer-like sites or transcription factor binding sites in HCT116 cells as determined by ENCODE ChIP-Seq data. Figure 3-figure supplement 3 lists the final target sites chosen for all 10 targeted regions.

Inspection of the ENCODE chromatin status in the 70 kb region surrounding each of these 10 target sites in HCT116 cells (Figure 3-figure supplement 4 and 5) confirms that the targeted lamina-associated/transcriptionally inactive regions have more heterochromatin characteristics as compared to the targeted speckle-associated/transcriptionally active regions which have more euchromatin characteristics, as expected (Ernst et al., 2011; Pombo and Dillon, 2015; Spector and Lamond, 2011). More specifically, all speckle-associated regions had enrichment of one or more of the chromatin marks (H3K4me3, H3K4me1, H3K27ac, H3K36me3) associated with active regions (Figure 3-figure supplement 4). All lamina-associated sites lacked these euchromatin marks and some had enrichment for repressive chromatin marks (H3K9me3, H3K27me3) (Figure 3-figure supplement 5). ChIP-Seq data for HCT116 cells were retrieved from the ENCODE Consortium.

To improve knock-in efficiency, we tried increasing the CRISPR plasmid to donor DNA fragment molar ratio to 2:1. We anticipated that maintaining a low transfected donor concentration would help minimize off-target integration; therefore, we did not increase the amount of donor DNA fragment. PCR-amplification of the donor DNA for all 10 regions yielded products showing single bands by gel electrophoresis (Figure 3A). Blasticidin selection for 5 days began one day after transfecting HCT116 cells with the donor DNA and CRISPR/Cas9 plasmid. PCR analysis of mixed pool cells after selection showed successful knock-ins for 9 out of the 10 targeted regions. (Figure 3B,C). Region 8 was the only locus for which we could not detect a knock-in PCR product. The sgRNA target site in region 8 overlaps with a region enriched in the heterochromatin marker H3K9me3 (Figure 3-figure supplement 5), which may have reduced knock-in efficiency at this location.

### Efficiency of knock-in at two euchromatin and two heterochromatin regions

We chose two heterochromatin (regions 9 and 10) and two euchromatin target sites (regions 3 (*HSP70*) and 5) for a comparison of knock-in efficiencies as a function of chromatin state. We carried out positive selection and clonal isolation simultaneously to shorten the overall transfection procedure. After clonal isolation, genotyping PCRs with Out-For and Out-Rev primers (Figure 3B) identified four clones from region 10 (10-13, 10-21, 10-22, 10-24), two clones from regions 3 (3-2, 3-9) and 5 (5-2,5-6), and one clone from region 9 (9-44) as potential correctly targeted, full length knock-ins (Figure 3D,E). Further analysis using the SpeI digestion-test of the knock-in bands revealed all these clones were correct except for 10-13 (Figure 3-figure supplement 6A). Analysis with both left and right junction PCRs further suggested most clones as correctly targeted and containing intact donor DNA (Figure 3-figure supplement 6B). However, unexpectedly we could not amplify the right junction PCR band from the 9-44 clone, suggesting a local loss of donor or flanking DNA sequence.

We sequenced the gDNA-insert junctions for all 8 remaining cell clones (3-2, 3-9, 5-2, 5-6, 9-44, 10-21, 10-22, 10-24) to determine whether the inserts had been integrated into the genome through HR (Figure 3-figure supplement 7). Interestingly, although all targeted insertions into euchromatic sites occurred through HR, all four targeted integrations into the gene-poor, heterochromatic regions 9 and 10 instead occurred through non-homologous end-joining (NHEJ). For these heterochromatin target regions, donor DNA was directly ligated after the expected cut site (Figure 3-figure supplement 7); moreover, various lengths of deletions of the linear donor DNA from their ends were observed in all clones (9-44, 10-21, 10-22, 10-24) which also explained why PCR-amplification of the right junction for clone 9-44 failed (Figure 3-figure supplement 7). Although integration occurred through NHEJ rather than HR, we counted these integrations as correctly targeted because this sequencing confirmed that the 48-mer TetO repeats were all intact. Our overall results therefore show a pronounced bias towards NHEJ-based integrations into lamina-associated, heterochromatin sites (Table 1).

**Table 1.** Summary of Genotyping Results

| Region Number | Expected Localization of the Region | Number of Clones Analyzed | Single Knock-in Clones | Double Knock-in Clones | Overall Knock-in Efficiency |
| --- | --- | --- | --- | --- | --- |
| 3 | Nuclear Speckle | 10 | 1/10 | 1/10 | 20% |
| 5 | Nuclear Speckle | 11 | 2/11 | 0/11 | 18% |
| 9 | Nuclear Lamina | 48 | 1/48 | 0/48 | 2% |
| 10 | Nuclear Lamina | 23 | 2/23 | 1/23 | 13% |

| Clone Number | Chromatin Status around Knock-in Site | Transcriptional Activity around Knock-in Site | 5' junction PCR | 3' junction PCR | Preferred integration mechanism |
| --- | --- | --- | --- | --- | --- |
| 3-2 | Euchromatin | High | + | + | HDR |
| 3-9 | Euchromatin | High | + | + | HDR |
| 5-2 | Euchromatin | High | + | + | HDR |
| 5-6 | Euchromatin | High | + | + | HDR |
| 9-44 | Heterochromatin | Low | + | - | NHEJ |
| 10-21 | Heterochromatin | Low | + | + | NHEJ |
| 10-22 | Heterochromatin | Low | + | + | NHEJ |
| 10-24 | Heterochromatin | Low | + | + | NHEJ |

Observed knock-in efficiencies were 20%, 18%, 2% and 13% for regions 3, 5, 9 and 10, respectively (Table 1), demonstrating that knock-in into gene-poor, heterochromatic sites is possible, albeit with lower efficiency. Moreover, the knock-in efficiency for region 3 may have slightly improved when CRISPR/Cas9 plasmid amount was increased. We observed two double knock-in clones: 3-2 and 10-22 for regions 3 and 10, respectively (Figure 3D, E and Table 1).

### 48-mer TetO repeat allows visualization of both euchromatin and heterochromatin sites

Next, we tested whether all four of the tagged loci could be visualized by the expression of TetR-EGFP. For more efficient labeling, a lentiviral TetR-EGFP-ires-PuroR vector was constructed in which *TetR-EGFP* and puromycin resistance (*PuroR*) genes are both expressed from the same promoter for more efficient positive selection. We used the weak F9 promoter to reduce the background coming from unbound TetR-EGFP. Clones 3-9, 5-2, 9-44 and 10-21 were used for visualizing regions 3, 5, 9 and 10, respectively. Cells were transduced with the lentivirus at low multiplicity of infection (MOI) and selected with puromycin for 3 days (Figure 4A). After selection, nearly 100% of cells expressed TetR-EGFP. Among the TetR-EGFP expressing cells, 40-70% of cells showed a distinct GFP spot visible above the background. In previous work using the lac operator / repressor system, detection sensitivity was a function of the lac repressor expression level, with low expression reducing the intensity over the lac operator repeats but high expression creating a GFP background level which masked the lac operator repeat signal (Robinett et al., 1996). All clones analyzed showed a distinct GFP spot in a fraction of the cell population (Figure 4B). Cells with a medium level of TetR-EGFP expression gave higher percentages of cells showing a distinct GFP spot. Subcloning TetR-EGFP stably expressing cells from the 3-9 original cell clone demonstrated that by selecting a subclone (3-9-5) with the appropriate level of TetR-EGFP expression, distinct GFP spots could be visualized in nearly 100% of cells.

**Figure 4:**
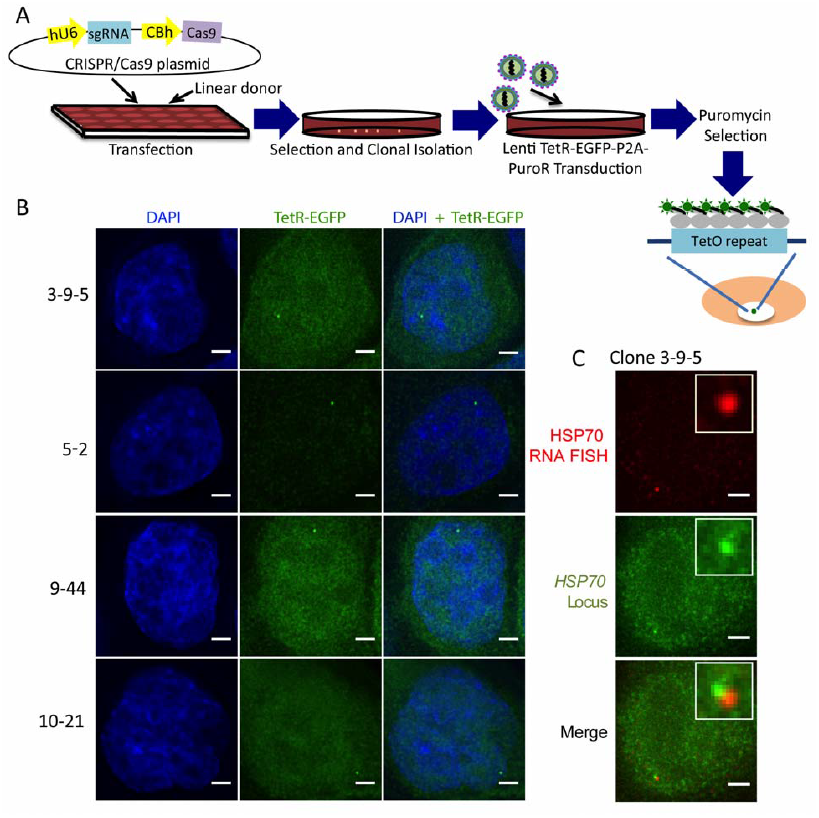
Visualization of 48-mer TetO-labeled endogenous loci by lentiviral expression of TetR-EGFP. **(A)** Overview of the workflow for the imaging of endogenous loci via knock-in of a 48-mer TetO repeat and expression of TetR-EGFP. **(B)** Imaging of 48-mer TetO-labeled regions 3, 5, 9 and 10 in clones 3-9-5, 5-2, 9-44 and 10-21, respectively by lentiviral expression of TetR-EGFP (green). Cells were fixed and stained with DAPI (blue). Single z sections are shown. Scale bar: 2 μ m. **(C)** Co-labeling of region 3 (*HSP70* locus) in clone 3-9-5 using TetO/TetR-EGFP system (Green) and RNA FISH (Red). All images are z projections. Scale bar: 2 μ m.

We used RNA FISH to confirm the specificity of the TetR-EGFP signal at the *HSP70* locus (region 3). The *HSP70* genes at the *HSP70* locus are heat-inducible. After heat-shock, RNA FISH revealed an accumulation of HSP70 nascent transcripts (red) immediately adjacent to the GFP spot tagging the *HSP70* locus (Figure 4C).

Using SR-SIM super-resolution light microscopy, we visualized GFP doublets in clones 3-9-5 and 10-21 as expected for sister chromatids following DNA replication (Figure 5A-C). The distances between these paired GFP spots ranged from 40-470 nm for region 3 and 80-830 nm for region 10 (Figure 5D). No cells with more than two GFP spots were observed (Figure 5E), and the distribution of distances showed no cases in which the distance between the two GFP spots exceeded 1 micron (Figure 5D). Clone 10-21 showed a lower percentage of cells with doublets versus a single spot as compared to the 3-9-5 clone, as predicted due to the expected late versus early DNA replication timing for regions 10 and 3, respectively (Figure 5E).

**Figure 5:**
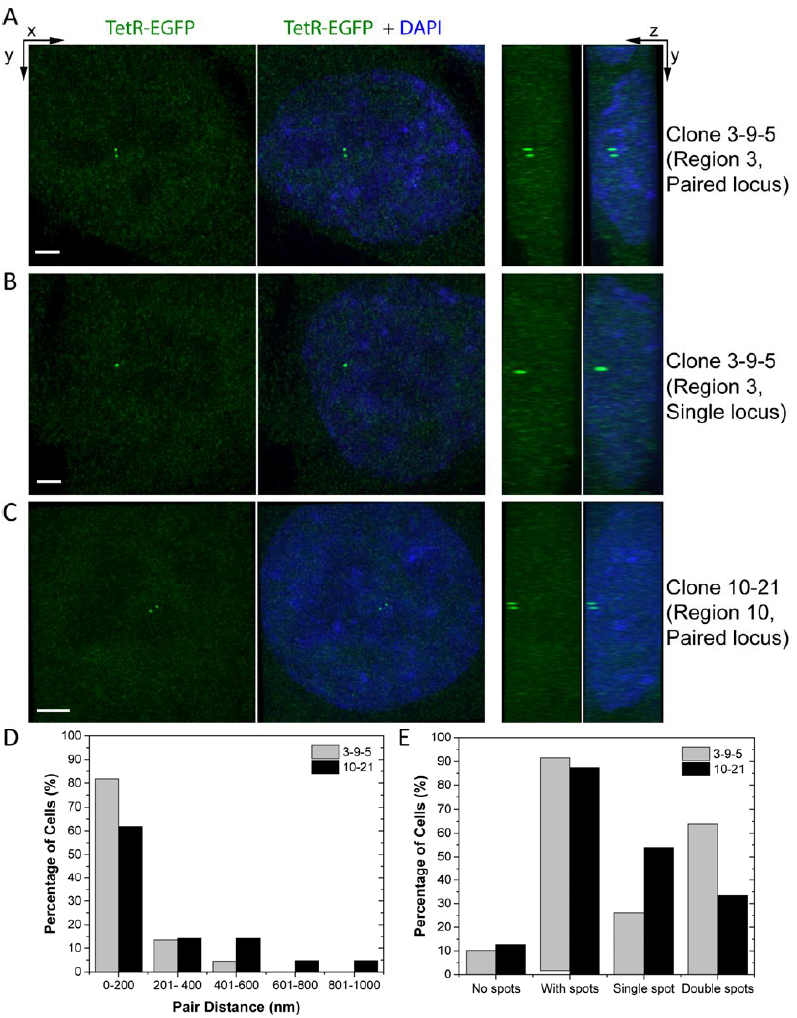
Analysis of the cells with paired loci. **(A)** 3D images of the 3-9-5 clone in SR-SIM microscope. 48-mer TetO-labeled region 3 was labeled by lentiviral TetR-EGFP (green) expression. Cells were fixed and stained with DAPI (blue). Images with EGFP only or EGFP+ DAPI are shown in XY (left 2 columns) and YZ (right 2 columns) projections. Here, images of a cell with paired spots **(A)** or a single spot **(B)** are shown. Scale bar: 0.7 μ m. **(C)** 3D images of the 10-21 clone in SR-SIM microscope. 48-mer TetO-labeled region 10 was labeled by lentiviral TetR-EGFP (green) expression. Cells were fixed and stained with DAPI (blue). Images of a cell with paired spots are shown. In this picture, the two spots are at the top of the nucleus. Scale bar: 1 μ m. **(D)** Distribution of the pair distance (in nm) between paired loci after DNA replication. Results for both region 3 (in clone 3-9-5, N=44 cells) and region 10 (in clone 10-21, N=21 cells) are shown. **(E)** Histograms of percentage of the cells with indicated number of spots observed per cell in 3-9-5 (N=69 cells) and 10-21 (N=63 cells) clones. % of cells with spots demonstrates the percentage of the total counted cells that had either a single spot or double spots.

The successful knock-in of the TetO repeat and its visualization in multiple euchromatic and heterochromatic chromatin regions demonstrates the general applicability of our method. We observed no extra GFP spots, which would have been derived from random integration of the donor DNA, in all 4 clones analyzed.

### Knock-in of the 48-mer TetO repeat does not perturb the intranuclear localization of the target chromosome locus

To verify that the insertion of the TetO repeat plus blasticidin selectable marker did not perturb the normal intranuclear targeting of the target chromosome locus, we compared the intranuclear localization of the euchromatic, speckle-associated *HSP70* and heterochromatic, lamin-associated *HBB* (regions 3 and 10, respectively) gene loci with or without the targeted donor DNA insertion. We compared the distance distribution of these chromosome loci from the nuclear lamina (*HBB*) and speckles (*HSP70*) in wt HCT116 cells versus for the knock-in alleles in clones 10-21 and 3-9-5, respectively (Figure 6). The loci in wt HCT116 were labeled by DNA-FISH using the corresponding BAC probes. No significant difference in distributions was observed (Figure 6C,D), indicating that insertion of the TetO repeat and blasticidin selectable marker did not perturb the normal intranuclear targeting of these chromosome loci.

**Figure 6:**
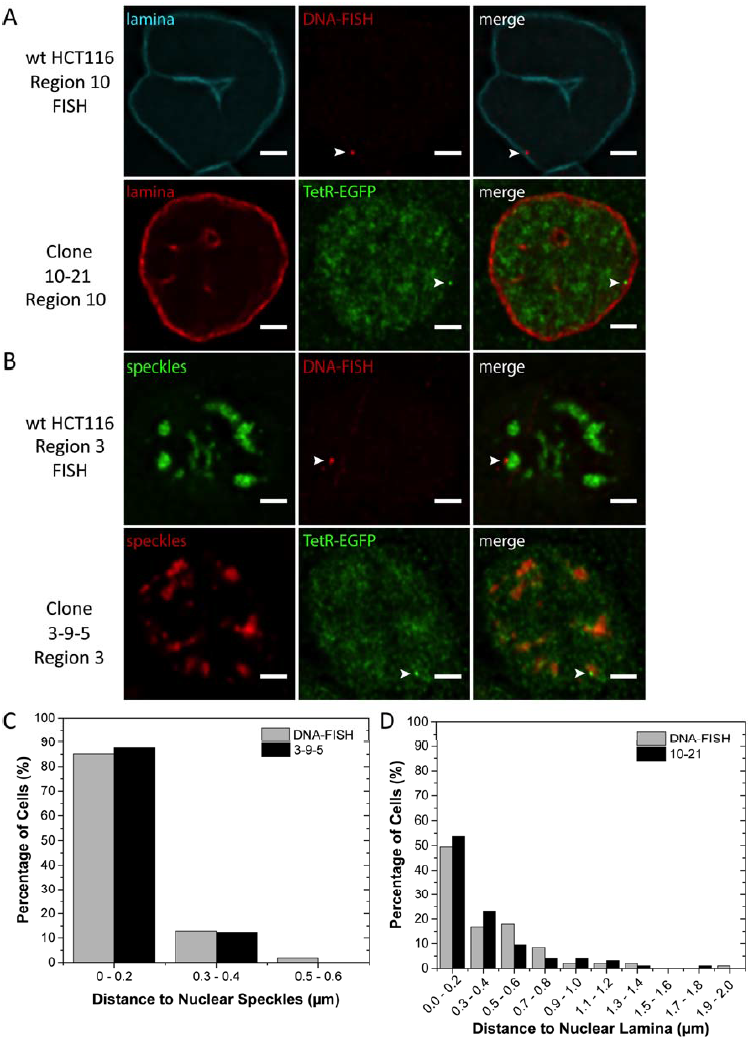
Controlling the effect of 48-mer TetO knock-in on the localization of endogenous loci. **(A)** Representative images of localization of the region 10 in wildtype (wt) HCT116 cells (top images), as determined by DNA-FISH (red), as well as in 10-21 48-mer TetO knock-in clone (bottom images), by expression of TetR-EGFP (green). Nuclear periphery was labeled with anti-laminB1/B2 antibody (shown in cyan for the top images and red for bottom images). The locus is pointed with an arrow. Single z sections are shown. Scale bar: 2 μ m. **(B)** Representative images of localization of the region 3 in wt HCT116 cells (top images), as determined by DNA-FISH (red), as well as in 3-9-5 48-mer TetO knock-in clone (bottom images), by expression of TetR-EGFP (green). Nuclear speckles were labeled with anti-SON antibody (shown in green for the top images and red for bottom images). The locus is pointed with an arrow. Single z sections are shown. Scale bar: 2 μ m. **(C)** Histogram of the distance between region 3 and the nuclear speckles in 3-9-5 knock-in clone (N= 97 cells) and wt HCT116 cells (N=101 cells). **(D)** Histogram of the distance between region 10 and the nuclear lamina in 10-21 knock-in clone (N=95 cells) and wt HCT116 cells (N=95 cells).

### Live cell imaging of the TetO tagged loci enables capturing the dynamics of loci with their associated nuclear compartments

Finally, we tested live cell imaging of the euchromatic and heterochromatic *HSP70* and β -globin loci (clones 3-9-5 and 10-21, respectively). First, we tested the stability and dynamics of the *HSP70* locus association with nuclear speckles. Measurement of distances to nuclear speckles in fixed cells (Figure 6C) had shown over 80% of loci within 0.2 microns of nuclear speckles, with nearly all remaining loci less than 0.5 microns from the speckles. Live cell imaging at 1 min intervals revealed that individual chromosome loci would occasionally move several tenths of a micron away from the nuclear speckle and back over short time intervals (Figure 7A,C and Movie 1). Thus the distance distribution observed in fixed cells was derived partially from small fluctuations in speckle distance over time at individual loci in live cells; however, overall speckle association remained stable over the 28 min observation periods. Similarly, live-cell imaging revealed oscillations of several tenths of microns of the *HBB* locus towards and away from the nuclear lamina over several min time intervals (Figure 7B,D and Movie 2), but with the locus remaining close to the nuclear lamina over the 28 min observation periods.

**Figure 7:**
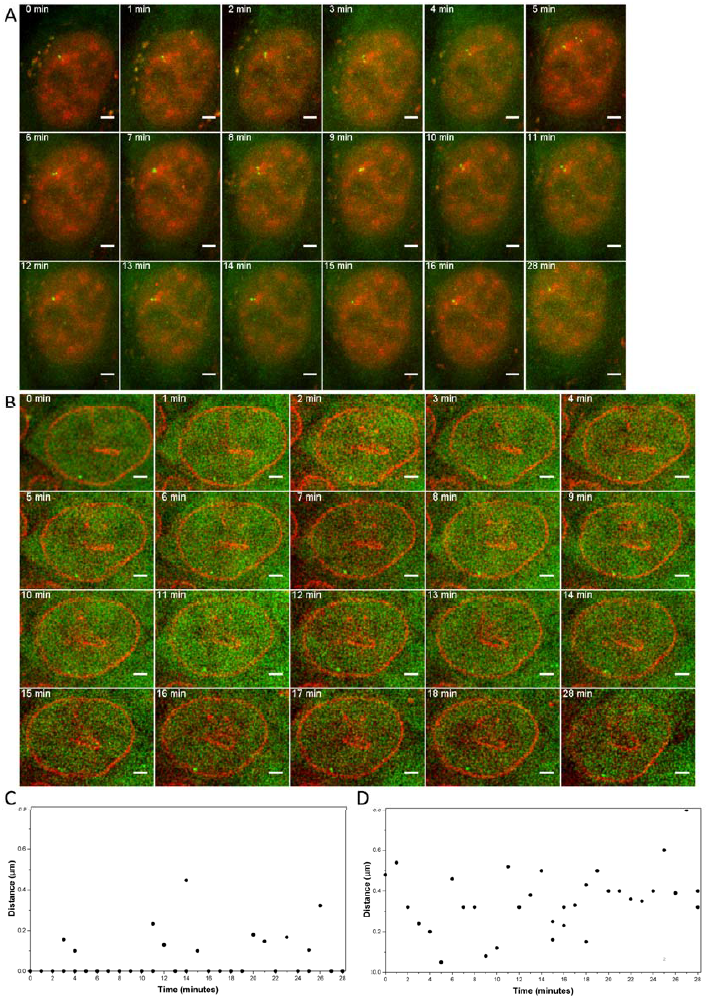
Dynamics of the nuclear speckle or lamina interactions. **(A)** Snapshots from the live cell imaging of region 3 (*HSP70* locus, images are from the 3-9-5 clone) and its interaction with nuclear speckles (red). Nuclear speckles were labeled by Magoh-mCherry expression. Interaction of the locus and nuclear speckles is stable over time. The movie was taken for 28 minutes with 1 minute time-lapse. All images are z projections. Scale bar: 2 μ m. Also see Movie 1. **(B)** Snapshots from the live cell imaging of region 10 (images are from the 10-21 clone) and its interaction with nuclear lamina (red). Nuclear lamina was labeled by mCherry-laminB1 expression. Interaction of the locus and the nuclear lamina is stable over time. The movie was taken for 28 minutes with 1 minute time-lapse. Single z sections are shown. Scale bar: 2 μ m. Also see Movie 2. **(C)** Histogram of the distance between region 3 and the nuclear speckles in 3-9-5 knock-in clone. **(D)** Histogram of the distance between region 10 and the nuclear lamina in 10-21 knock-in clone. For **(C)** and **(D)**, when two spots were observed, both of them were plotted.

## Discussion

Here we demonstrated high efficiency knock-in of an optimized 48-mer TetO repeat for labeling and live cell tracking of various chromosome endogenous loci of interest. Our targeted insertion method was effective at both euchromatic and heterochromatic chromosome loci with varying transcriptional and chromatin status. Importantly, our tagging method did not perturb the normal intranuclear localization of the unmodified *HSP70* and β -globin chromosome loci. As presented, we showed the usefulness of this TetO tagging method for cytological analysis in both fixed and live cells. We anticipate further development of this tagging technology, for instance by using split-GFP (Cabantous et al., 2005) or SunTag imaging approaches (Tanenbaum et al., 2014), will provide increased signal-to-background and noise ratios allowing longer-term imaging with reduced light exposure and phototoxicity.

Random integration of exogenous DNA has always been a concern in knock-in strategies; however, our imaging of the targeted clones revealed only the number of GFP-tagged loci predicted by our genotyping and FISH analysis. Therefore the frequency of nonspecific integration of donor DNA is low using our experimental targeting conditions. The observed low frequency of off-target integrations may be due to our exclusion of repetitive and low-complexity sequences from the short homology arms used in our approach. The high repetitive DNA content of mammalian genomes makes exclusion of such repetitive and low-complexity sequences difficult using longer homology arms. Thus, our use of short homology arms might actually have promoted a high frequency of correctly targeted integrations. At the same time, it is likely that the low amount of donor DNA used for transfection helped minimize random integrations. Similar targeting conditions might be beneficial for other applications of homology-directed genome modifications such as gene correction in various genetic diseases.

Our labeling strategy to visualize non-repetitive regions in the genome requires minimal modification of the genome: the only two components that need to be integrated into the genome are the 48-mer TetO repeat and the TetR-EGFP expression cassette. CRISPR/Cas9 only needs to be transiently expressed and its integration into the genome is unnecessary. This is an advantage over previous CRISPR/Cas9-based imaging techniques that required integration of at least 26 sgRNAs, each from a separate lentivirus, per cell (Chen et al., 2013). While our manuscript was in preparation, an improved CRISPR/Cas9-based method for visualizing non-repetitive regions was published (Qin et al., 2017). In this paper, sgRNAs were tagged with multiple MS2 repeats and visualized by expression of MCP-FP fusion proteins. Although Qin et al. was able to increase signal-to-noise ratio significantly for visualization of non-repetitive sites, the system still requires integration of a minimum of 8 different sgRNAs for visualization using conventional fluorescence microscopy. Moreover, stable expression of both MCP-FP and dCas9 from separate lentiviruses is also necessary, requiring a total of 10 lentiviruses that need to be integrated per cell. Our method provides a complementary alternative to TALE and dCas9 based visualization methods requiring fewer components to be transfected into the target cells. We anticipate that using different repetitive sequences-for instance lacO repeats or repeats corresponding to binding targets of single TALE proteins or sgRNAs-will allow extension of our methodology to the tagging of multiple endogenous chromosome loci. What remains to be determined are the relative advantages and disadvantages of each of these different chromosome tagging systems. For example, although CRISPR/dCas9 is now being widely adopted for live-cell imaging, we still do not know whether the R-loops created by the CRISPR/dCas9 system will perturb cell physiology, as do naturally occurring R-loops (Skourti-Stathaki and Proudfoot, 2014). Future comparisons by live-cell imaging of the same chromosome locus tagged with different systems (TALEs, CRISPR/dCas9, or TetO repeats) should prove useful in addressing these questions.

Finally, our results are also informative with respect to understanding the relationship between chromatin state and transcriptional activity and the choice of DSB repair mechanism used at different chromosome loci. A previous publication suggested heterochromatin and transcriptionally silent sites avoid HR (Aymard et al., 2014). Our results showing targeted insertion at heterochromatin targets by NHEJ rather than HR are consistent with this previous finding. We also note that the one target locus for which we saw no PCR product indicative of targeted insertion showed a noticeable enrichment of the H3K9me3 heterochromatin mark. Further analysis should clarify better which histone marks and subnuclear compartmentalization effect knock-in efficiency and/or DSB repair pathway choice, which would lead to improved rules for sgRNA design.

## Acknowledgements

This work was supported by National Institutes of Health grants 1U54DK107965 (H.Z., J.M. and A.S.B.) and R01GM058460 (A.S.B).

## Materials and Methods

### Assembly of TetO Repeats

First, an initial cassette of three Tet operators (19 bp) and four random sequences (10 bp) separating the TetO sequences was designed. For that purpose, a library of two types of oligonucleotides (82 bp and 83 bp) were synthesized (Integrated DNA Technologies (IDT), Coralville, IA):

1^st^ oligo: 5’-CACAGGAAACAGCTATGACCCCTAGGNNNNNNNNNN<u>TCCCTATCAGTGATAGAGA</u>NNNNN NNNNN<u>TCCCTATCAGTGATAGA</u> -3’

2^nd^ oligo: 5’-<u>GA</u>NNNNNNNNNN<u>TCCCTATCAGTGATAGAGA</u>NNNNNNNNNNACTAGTTAGGATGAAGGG ATCCGTTGTAAAACGACGGCCAGT -3’

Random 10 bp sequences are indicated with N, and sequences for TetO is underlined. 10 bp random sequences contained only A, G, and T (avoided C to prevent forming of CG sites), randomly mixed in the ratio of 25:50:25. Ratio of nucleotides was chosen to mimic an average GC content in the genome. Three restriction sites (BamHI, SpeI and StyI, shown in bold) were also included in the design for subsequent steps to synthesize TetO multimers. Two of these enzymes have compatible sticky ends (SpeI, StyI) and after ligation it creates a site that cannot be recut again with either enzyme (Robinett et al., 1996). None of the restriction enzymes contained CG in the sequence and they were unique cutters in the backbone of the plasmid used for cloning. We ordered a library of the 1^st^ and the 2^nd^ oligos with various 10 bp random sequences.

To be able to ligate two oligo pieces together and get a library of 3mer of TetO repeats (165 bp), second oligos were phosphorylated at the 5’end. Ligation of two ssDNAs was performed by T4 RNA ligase (New England Biolabs (NEB), Ipswich, MA) according to manufacturer’s instructions. Briefly, 1x reaction buffer, 25% PEG, 1 mM ATP, 1 mM hexamine cobalt chloride, 10 units T4 RNA ligase, and 0.5 μ M of each pool of ssDNA oligonucleotides were mixed together and incubated at 22°C for 16 hours, followed by heat inactivation for 15 minutes at 65°C. The resulting ligated 165 bp product (Figure 1B) was checked on 3% agarose gel.

After ligation of the two oligonucleotides, synthesis of 2^nd^ DNA strand followed. Complementary strand was synthetized by PCR with high-fidelity *Pfu* Turbo DNA Polymerase (Agilent Technologies, Santa Clara, CA) using primers flanking the initial cassette (highlighted in yellow within the sequences for oligos). Reaction was set up according to manufacturer’s instructions with 0.5 μ l of ligation product serving as a template and the PCR was ran for 15 cycles. The same polymerase and primers were used for 2^nd^ PCR with 0.5 μ l of the 1^st^ PCR product used as a template. Correct size of the final product was confirmed by gel electrophoresis.

Our goal was to create 96-mer of TetO repeats, which meant to obtain 32 unique TetO 3mers as the building blocks. We used a low copy number plasmid pSP2 for cloning to reduce risk of unwanted recombination and ensure the stability of the carried repeats (Carpenter and Belmont, 2004). Initial insertion of TetO 3mer into the pSP2 vector was done in following steps (Figure 1C):

<u>*Step 1 and 2*:</u> Digestion of the pSP2 vector and the insert (PCR Product) with StyI and BamHI (NEB).

<u>*Step 3*:</u> Gel purification of the vector and the insert using Gel Extraction Kit (Qiagen, Valencia, CA).

<u>*Step 4*:</u> Ligation of the insert with the vector using FastLink DNA Ligation Kit (Epicentre, Madison, WI) according to manufacturer’s instructions. 75 ng of vector DNA was used for most of the ligation reactions. Ligation mixture was incubated for 30 minutes at room temperature (RT) and enzyme was heat inactivated for 15 minutes at 70°C.

After insertion of 3mer TetO into the pSP2 vector, we used special Library Efficiency^^®^^ DH5α ™ Competent Cells (LECC) (Invitrogen, Carlsbad, CA) for transformation of the ligation product. High transformation efficiency of these cells ensured obtaining a large number of clones with different random sequences separating TetO repeats and thus allowed us to obtain a library of TetO 3mers. According to manufacturer’s instructions, 2.5 μ l of the ligation product was transformed to 100 μ l of LECC cells by heat shock for 45 seconds at 42°C. After recovery in 900 μ l of SOC medium for 1 hour, cells were seeded on 6 plates with proper antibiotic selection (Ampicillin, 100 μ g/ml). All colonies were scraped from the plates using a plastic cell scraper and homogenized in resuspension buffer in 1.5 ml eppendorf tube. Few μ l of the cell suspension was inoculated into 1 ml of LB medium, incubated for 6 hours at 37°C and frozen in 25% glycerol (this served as a library of TetO 3mers). The rest of the homogenized bacterial culture was used for DNA isolation using Plasmid Mini Kit (Qiagen) with a yield of around 5 μ g.

To create a library of TetO 6mers, 2 μ g of DNA was used for preparation of the insert and the backbone vector (both now containing TetO 3mer). DNA serving as a backbone vector was cleaved with SpeI and BamHI, and DNA to obtain the insert from was digested with StyI and BamHI (NEB; Figure 1D). Both the vector and the insert were gel purified and then ligated, similar to the protocol for creating 3mer TetO library. Transformation into LECC cells resulted in a library of TetO 6mers, which was frozen the same way as TetO 3mer library (described above). Additionally, 19 individual colonies containing correct 6mers were isolated.

After assembly of 6mer TetO repeats, we switched to a safer approach to assemble higher multimers step by step. We wanted to be sure that all clones contained correct sequence of TetO repeats and that all random 10bp parts are unique. The 19 isolated 6mer plasmids had random 10 bp parts that were all unique. Eight ligation reactions were performed to get eight different 12mers. Pairs of 6mer plasmids (one to serve as vector backbone and the other to digest-out the insert, Figure 1D) were used for ligation. The same process was repeated in four ligation reactions to obtain four different 24mers.

Since there was no need for high efficiency competent cells anymore, we used standard DH5α competent cells for transformation of the ligation products. Presence and correct size of TetO cassette was checked by PCR or restriction digestion with StyI and BamHI. All four 24mers were sequenced and all had correct TetO sequence. After final ligation of two 48-mers we got pSP2 plasmid containing TetO 96-mer with size around 3.3 bp.

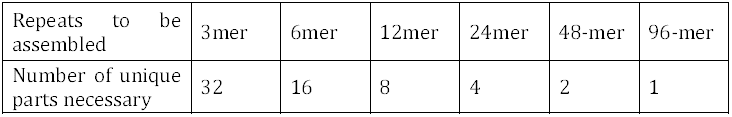

| Repeats to be assembled | 3mer | 6mer | 12mer | 24mer | 48-mer | 96-mer |
| --- | --- | --- | --- | --- | --- | --- |
| Number of unique parts necessary | 32 | 16 | 8 | 4 | 2 | 1 |

## Plasmid Construction

To create a plasmid to serve as a template for linear donor DNA PCRs, pSP2 48-mer TetO and pSP2 96-mer TetO plasmids were further modified. All the following modifications were done in parallel to both pSP2 48-mer TetO and pSP2 96-mer TetO plasmids. First, a gBlock containing a multiple cloning site, flanked by loxP sites (shown in bold below), was synthesized (IDT). The gBlock also had the sequence for rop protein (in italic and underlined), which maintains plasmids at low copy number. The gBlock had SpeI and SapI sites at 5’ and 3’ ends, respectively (underlined in the sequence below). The sequence for gBlock was as followed:

5’-GGCAGCTGTTG<u>ACTAGT</u>ATAACTTCGTATAATGTATGCTATACGAAGTTATCATATGGGCT AGCTAGCTAGGAATTCGAAGCTTGAGCTCGAGATCTGAGTGGATCCTAGTTCGAAATAACTTCGTATAATGTATGCTATACGAAGTTATCTCGAGAGCGCTGCTAGCTTAAGGTACCATCGATAGAT CTGGGCCC<u>GTGACCAAACAGGAAAAAACCGCCCTTAACATGGCCCGCTTTATCAGAAGCCAGACATTA</u> <u>ACGCTTCTGGAGAAACTCAACGAGCTGGACGCGGATGAACAGGCAGACATCTGTGAATCGCTTCACGA</u> <u>CCACGCTGATGAGCTTTACCGCAGCTGCCTCGCGCGTTTCGGTGATGACGGTGAAAACCTCTGA</u>CTTA AGCCTGAG<u>GCTCTTC</u>CGCTTTG-3’

The gBlock was cloned through SpeI and SapI sites into the pSP2 48-mer/96-mer TetO plasmids. As a next step, EFS promoter from lentiCRISPR v2 plasmid (A gift from Feng Zhang, Addgene plasmid # 52961) was PCR-amplified with the following forward and reverse primers that introduced NdeI and HindIII sites (underlined), respectively: For 5’-TTTG<u>CATATG</u>GCTAGGTCTTGAAAGGAGTGGG-3’ and Rev 5’-GTTT<u>AAGCTT</u>CCTGTGTTCTGG CGGCAAA-3’. The PCR product was cloned into pSP2 48-mer/96-mer TetO loxP plasmids through NdeI and HindIII sites (between the 2 loxP sites). The resulting plasmids were further modified by cloning a blasticidin resistance (BlaR) gene and SV40 polyA sequence downstream the EFS promoter through HindIII and BstBI sites. For that purpose, BlaR gene from pF9-TALE-EGFP-Blast plasmid was PCR-amplified with the following forward and reverse primers that introduced HindIII-Kozak sequence and BstBI sequence (restriction sites are underlined, Kozak sequence is in bold), respectively: For 5’-TTTG<u>AAGCTT</u>GCCACCATGGCCAAGCCTTTGTCTC-3’ and Rev 5’-GTTGTT<u>TTCGAA</u>ACATGTGA TCCAGACATGATAAGAT AC-3’.

To create lenti F9-TetR-EGFP-IRES-PuroR plasmid in which both TetR-EGFP and PuroR genes are expressed from the F9 promoter, F9-TetR-EGFP was PCR-amplified from the p3’SS TetR-EGFP vector with following forward and reverse primers that introduced SpeI and PshAI sites (underlined), respectively: For 5’-TTGTTG<u>ACTAGT</u>ACGCGTATAGATCTGG ATCCCG-3’ and Rev 5’-CAACAA<u>GACCAGAGTC</u>TTACT TGTACAGCTCGTCCATGCC-3’. The PCR product was digested and cloned into Cre-IRES-PuroR vector (A gift from Darrell Kotton, Addgene plasmid # 30205) through SpeI and PshAI sites, upstream of IRES sequence, using traditional restriction digestion/ligation protocols.

To create mCherry-LaminB1, GFP in the GFP-laminB1 plasmid (a gift from Veena K Parnaik) was replaced with mCherry. Sequence for mCherry was digested out from the mCherry-PCNA-19-SV40NLS-4 vector (A gift from Michael Davidson, Addgene plasmid # 55117) with AgeI and BsrGI enzymes and cloned into GFP-laminB1 plasmid through the AgeI and BsrGI sites, downstream of CMV promoter, using traditional restriction digestion/ligation protocols. The final sequence was confirmed by Sanger sequencing (Genewiz, South Plainfield, NJ) for all of the plasmids.

### sgRNA and Homology Arm Design

BACs containing human DNA inserts within selected regions were identified (Figure 3-figure supplement 2) and the DNA sequences corresponding to these inserts (Region 1-10) were retrieved from UCSC Genome Browser (Kent et al., 2002). Transcriptional status of these regions were determined based on the ENCODE Caltech RNA-Seq data for HCT116 and K562 cells (The ENCODE Project Consortium, 2012), which we displayed on the UCSC Genome Browser (Rosenbloom et al., 2013). RNA-Seq data for HCT116 (UCSC accession number: wgEncodeEH001425) and K562 (UCSC accession number: wgEncodeEH000124) cells were created by Wold lab as part of the ENCODE Project (The ENCODE Project Consortium, 2012).

To design 20 nt guide sequences of sgRNAs for knock-in into each region of interest, DNA sequences of the sub-regions without any exons were retrieved from the UCSC Genome Browser. UCSC Genome Browser on Human Feb. 2009 (GRCh37/hg19) Assembly was used for all the analysis in this paper. Within the sub-regions, repetitive and low complexity DNA sequences were annotated by RepeatMasker (Smit et al., 1996-2010; <u>http://www.repeatmasker.org</u>) in the UCSC Genome Browser and we downloaded sequences with masked repeats. The unmasked regions were given to CRISPRdirect web tool (Naito et al., 2015) using a custom Application Program Interface (API) developed in Python programming language. The guide sequences within the unmasked sequences for each region were combined and sorted to find the most efficient and specific sgRNA sequence. All guide sequences that could result in difficulty in DNA synthesis (more than six consecutive A’s, C’s and G’s), sgRNA transcription (more than four consecutive T’s) and DNA assembly (BpiI recognition site) were removed from the list. The remaining guide sequences were then sorted based on GC content, specificity, and efficiency using the scoring criteria described below:

|  | Criterion | Weight (W) | Condition | Multiplier (M) |
| --- | --- | --- | --- | --- |
| Efficiency score | GC number | $W_1=1/2$ | 7 to 15 (including 7 and 15) | $M_1=1$ |
| | | | Less than 7 or more than 15 | $M_1=0$ |
| | Composition of the last four nucleotides | $W_2=1/2$ | | $M_2=0.25 \times (\#G) + 0.2 \times (\#A) + 0.15 \times (\#C)$ |
| Specificity score | $1/(\text{hit\_12mer})^2$ | | | |
| Total score | $100 * (M_1W_1 + M_2W_2) / (\text{hit\_12mer})^2$ | | | |

Where the efficiency score comprises of GC content of the guide sequence as well as the composition of last four nucleotides next to PAM sequence. The weight for the effect of each nucleotide on the efficiency is obtained from a previously published work (Wang et al., 2014). Specificity score of the guide sequence was evaluated based on the number of times the 12 nt sequence immediately upstream of the PAM is repeated in the genome, giving a very low score to guide sequences with non-specific 12 nt seed sequence.

For minimum disruption to the genome function, candidate sgRNAs with high scores (67 or more) were further evaluated in terms of their alignment with regulatory sites. UCSC Genome Browser (Kent et al., 2002) was used to search for possible transcription factor binding sites (TFBS). We used Transcription Factor ChIP-seq (161 factors) from ENCODE with Factorbook Motifs track to afind potential binding sites (Kent et al., 2002; Rosenbloom et al., 2013). We also retrieved enhancer-like sequences (http://zlab-annotations.umassmed.edu/enhancers/) and promoter-like sequences (http://zlab-annotations.umassmed.edu/promoters/) in HCT116 cells (Rosenbloom et al., 2013; The ENCODE Project Consortium, 2012). sgRNA target sites that align with any of these three regulatory sites were excluded.

As a final step, sgRNAs with predicted DSB sites (3 nt upstream of PAM) closer than 50 nt to the masked regions were omitted to provide a unique sequence for the homology arms. If an sgRNA with a good score could not be found within the regions in Figure 3-figure supplement 2, then we also screened the sequences that were around 50 kb upstream or downstream of the region. A list of selected sgRNA target sites can be found in Figure 3-figure supplement 2.

DNA sequences of 50 bp upstream and downstream of the predicted DSB site were retrieved and used as the homology arms for the HR. The homology arms were designed to exclude PAM sequence and also the first 3-4 nt upstream of the expected cut site (See Figure 3-figure supplement 3). Point mutations were introduced to the PAM if the homology arm included the PAM sequence (as for Region 3). Homology arms were further optimized to minimize secondary structures. A demonstration of the homology arms selected for each region can be found in Figure 3-figure supplement 3.

### sgRNA Cloning

Px330A-1×3 plasmid, a gift from Takashi Yamamoto (Sakuma et al., 2014), was used to express both sgRNA and Cas9 from the same plasmid. Oligonucleotides used for cloning the guide sequence into pX330A-1×3 plasmid can be found below. If the guide sequence did not start with a G, we added a G to the 5’ of the 20 nt sequence to improve transcription from the U6 promoter. Complementary oligonucleotides were ordered from IDT, annealed, phosphorylated and cloned into pX330A-1×3 plasmid through the BpiI sites by following a previously published protocol for the pX330 plasmid (Ran et al., 2013).

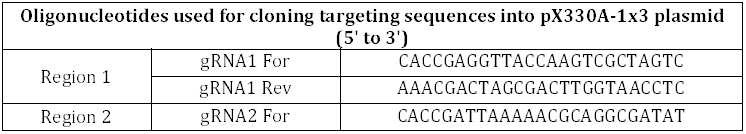
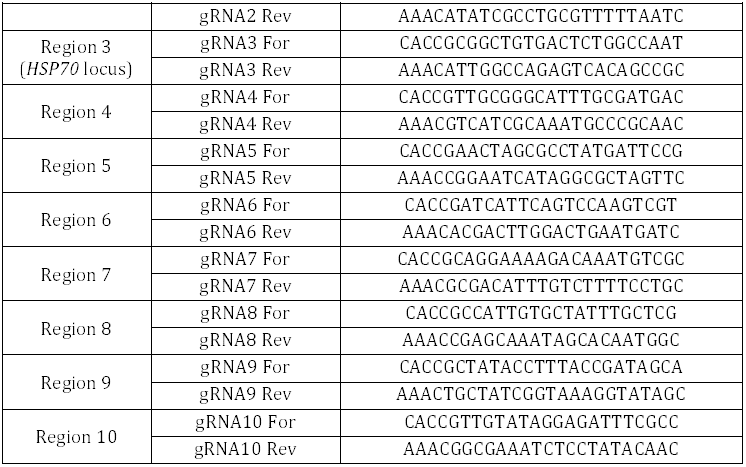

### PCR for Creating Linear Donor DNA

Sequences of the primers used for creating linear donor DNAs for targeting each of the 10 regions can be found in the table below. For the initial comparison between 48-mer and 96-mer TetO knock-in at the *HSP70* locus (Figure 2), the donor DNA was created with the primers “old donor3 For” and “donor3 Rev”. To create linear donor DNA for 10 regions in the later knock-in experiments (Figure 3), the part of the forward primer that anneals to the template was changed to add 66 nt more to the 5’ end of the insert. We aimed easier left junction sequencing by this design. Thus, for the later experiments to knock-in into region 3, donor DNA was created with the primers “donor3 For” and “donor3 Rev”. The parts of the primers that anneal to the template plasmid are shown in bold in the table below.

Primers were ordered from IDT and they contain phosphorothioate bonds between the last 3 nucleotides at the 5’ end of the primers. When transfected into mammalian cells, linear DNA can be attacked by cellular nucleases and be degraded, lowering the transfection efficiency. Thus, we introduced phosphorothioate bonds to overcome this problem (Orlando et al., 2010). The location of phosphorothioate bonds are shown with “* “in the table below. Forward primers contain a 23 nt sequence annealing to the template plasmid and upstream of it is the 50 nt homology sequence. In reverse primers, the sequence annealing to template is 21 nt. Either pSP2-48-merTetO-EFS-BlaR or pSP2-96-merTetO-EFS-BlaR plasmid was used as a PCR template, depending on the desired final donor DNA. PCRs were performed using Q5 High Fidelity DNA Polymerase (NEB), using manufacturer’s protocol with the following changes to the PCR cycle: 98°C for 3 minutes; followed by 30 cycles of 98°C for 30 seconds, 70°C for 15 seconds and 72°C for 3 minutes. The final extension was 72°C for 6 minutes. The donor DNA PCRs were gel-purified with Gel Extraction Kit (Qiagen) and eluted with ddH2O.

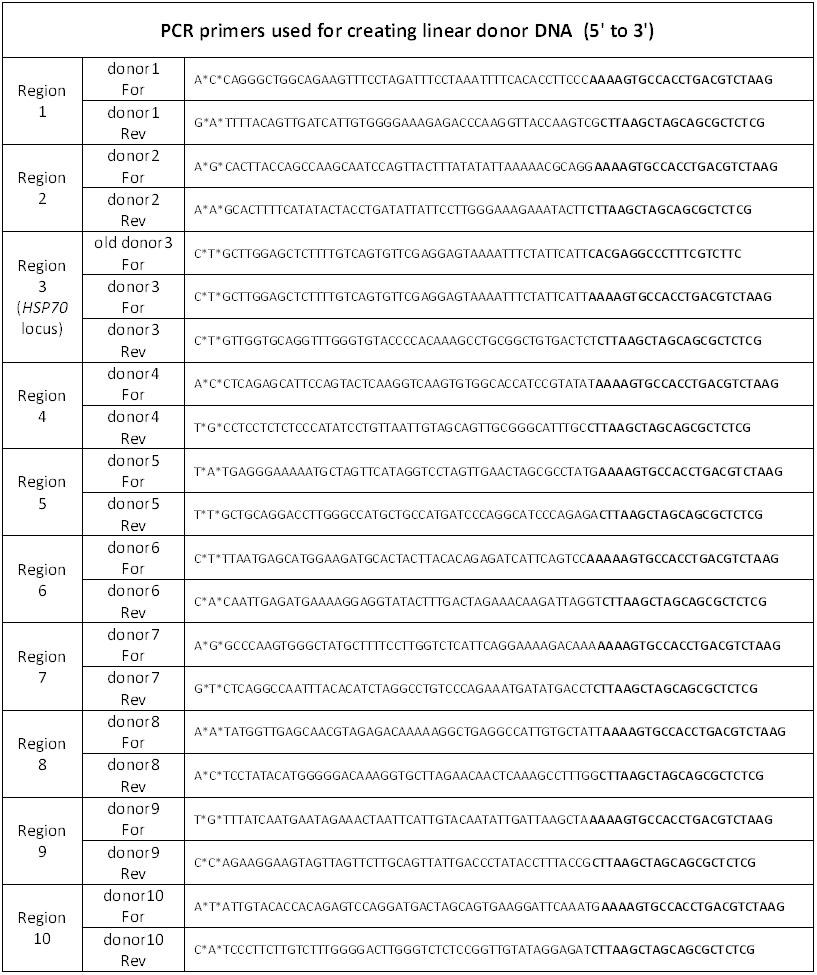

### ChIP-Seq data analysis

We analyzed six different histone modification ChIP-Seq data that were available for HCT116 cell line in the ENCODE database (The ENCODE Project Consortium, 2012).

Histone modification ChIP-Seq data for HCT116 cell line were downloaded from ENCODE project portal (<u>https://www.encodeproject.org/).</u> H3K4me3 ChIP-seq data (ENCODE accession number ENCSR000DTQ) were produced by the Stamatoyannopoulos laboratory; H3K4me1 (ENCODE accession number ENCSR000EUS), H3K27Ac (ENCODE accession number ENCSR000EUT), H3K9me3 (ENCODE accession number ENCSR000FCP) and H3K36me3 ChIP-seq data (ENCODE accession number ENCSR000FCQ) were produced by the Farnham laboratory; and H3K27me3 ChIP-seq data (ENCODE accession number ENCSR000BDB) were produced by Bernstein laboratory as part of the ENCODE Consortium. Note that for ChIP-seq data with biological replicates for the same histone mark, we merged the ChIP-seq BAM files of the replicates.

ChIP-Seq peaks were called with MACS2 (<u>https://github.com/taoliu/MACS</u>). MACS2 parameters: macs2 callpeak -t signal.bam -c control.bam -f BAM -g hs -nomodel -shiftsize 100 -n output_name -B -q 0.01). ChIP-Seq peaks are classified into narrow peaks (H3K4me1, H3K4me3, H3K27ac) and broad peaks (H3K27me3, H3K36me3, H3K9me3) based on signal enrichment patterns. The results were shown on the UCSC Genome Browser (Rosenbloom et al., 2013). Below is a summary of the ENCODE ChIP-Seq data used:

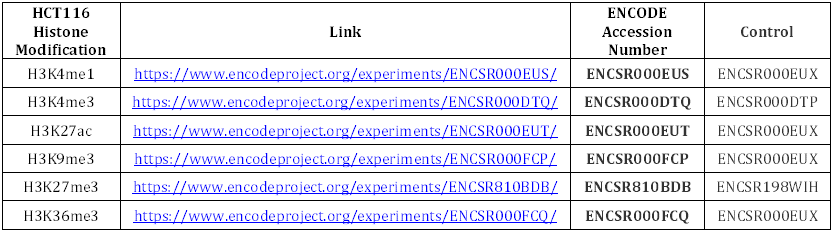

| HCT116 Histone Modification | Link | ENCODE Accession Number | Control |
| --- | --- | --- | --- |
| H3K4me1 | https://www.encodeproject.org/experiments/ENCSR000EUS/ | ENCSR000EUS | ENCSR000EUX |
| H3K4me3 | https://www.encodeproject.org/experiments/ENCSR000DTQ/ | ENCSR000DTQ | ENCSR000DTP |
| H3K27ac | https://www.encodeproject.org/experiments/ENCSR000EUT/ | ENCSR000EUT | ENCSR000EUX |
| H3K9me3 | https://www.encodeproject.org/experiments/ENCSR000FCP/ | ENCSR000FCP | ENCSR000EUX |
| H3K27me3 | https://www.encodeproject.org/experiments/ENCSR810BDB/ | ENCSR810BDB | ENCSR198WIH |
| H3K36me3 | https://www.encodeproject.org/experiments/ENCSR000FCQ/ | ENCSR000FCQ | ENCSR000EUX |

| Controls (ENCODE Accession Number) | Link |
| --- | --- |
| ENCSR000EUX | https://www.encodeproject.org/experiments/ENCSR000EUX/ |
| ENCSR000DTP | https://www.encodeproject.org/experiments/ENCSR000DTP/ |
| ENCSR198WIH | https://www.encodeproject.org/experiments/ENCSR198WIH/ |

### Cell Culture, Transfection and Clonal Isolation

Human embryonic kidney (HEK) cell line HEK293T was maintained in Dulbecco’s modified Eagle’s Medium (DMEM) (Corning Life Sciences, Tewksbury, MA) supplemented with 10% heat inactivated FBS (Life Technologies, Carlsbad, CA) at 37°C and 5% CO2 incubation. Cells were grown on BioCoat Collagen-I coated plates (Corning Life Sciences).

HCT116 cells were a gift from Mark S. Kuhlenschmidt and were cultured in McCoy’s 5A medium without phenol red (UIUC Cell Media Facility), supplemented with 10% tetracycline-free FBS (Sigma-Aldrich, St. Louis, MO). Cells were grown at 37°C in a humidified 5% CO2 incubator. BioCoat Collagen-I coated plates (Corning Life Sciences) were used for better attachment of the HCT116 cells. To perform transfections for knock-in of the TetO donors, Fugene HD (Promega, Madison, WI) was used, following manufacturer’s protocol. For the initial transfections to test knock-in efficiencies of the 48-mer and 96-mer TetO EFS-BlaR donors at the *HSP70* locus, cells were grown in 6-well plates until ~ 70% confluency. We used 1:1 CRISPR/Cas9 to donor molar ratio and we transfected 1 μ g of CRISPR/Cas9 plasmid. Accordingly, 0.3 μ g of 48-mer TetO EFS-BlaR donor and 0.5 μ g of 96-mer TetO EFS-BlaR donor were used. We started blasticidin (10 μ g/ml) selection 1 day after transfection. 7 days after blasticidin selection, clonal isolation was started by limiting dilution in 96-well plate. For the later transfections to knock-in 48-mer TetO EFS-BlaR donor into 10 different regions, including *HSP70* locus (region 3) as a positive control, we used 2:1 CRISPR/Cas9 to donor molar ratio. Before transfection, cells were grown in 24-well plates until 40-50% confluency. We transfected 500 ng CRISPR/Cas9 plasmid and calculated the necessary linear donor DNA amount accordingly (83 ng). 1 day after transfection, cells were plated onto 100 mm plates (together with 10 μ g/ml blasticidin) at limited dilution for growth of isolated colonies. Clonal isolation was performed following a previously published protocol (Strukov and Belmont, 2008).

3-9-5 clone was transfected with Magoh-mCherry plasmid (a gift from K. V. Prasanth) using Lipofectamine 2000 (Thermo Fisher Scientific, Waltham, MA), following manufacturer’s protocol. Briefly, cells were grown in T-25 flasks until 60-70% confluency and transfected with 3 μ g plasmid. Selection with G418 was started 2 days after transfection and cells were analyzed once the selection was completed.

10-21 clone was transfected with mCherry-laminB1 plasmid using Fugene HD (Promega), following manufacturer’s instructions. Briefly, cells were grown in 6-well plates until around 50% confluency and transfected with 1 μ g plasmid. Cells were analyzed 2 days after transfection. Same transfection protocol was followed for the transfection of the p3’SS TetR-EGFP plasmid into the 48-15 clone.

### Genotyping and Sequencing

Once the clones grown in 24-well plates reached ~ 80% confluency, half of the cells were collected and gDNA was extracted using QuickExtract DNA Extraction Solution (Epicentre). Briefly, 50 μ l QuickExtract solution was added per cell pellet, vortexed until a homogenous solution was obtained and then the mixture was incubated at 65°C for 15 minutes, followed by 98°C for 10 minutes. 4 μ l from the gDNA was used for genotyping PCRs. All the genotyping PCRs were performed using Q5 High Fidelity DNA Polymerase (NEB) in a 15-μ l reaction, using manufacturer’s protocol. We used the following PCR cycle: 98°C for 3 minutes; followed by 35 cycles of 98°C for 30 seconds, 68°C for 15 seconds and 72°C for 3 minutes (or 5 minutes for the 96-mer TetO knock-in clones). The final extension was at 72° for 6 minutes. Initially, PCR was performed using the out-forward and out-reverse genotyping primers for each region, listed in the table below. These primers both anneal to the gDNA outside of the homology arms. For the clones which had a band size similar to the size expected from the knock-in allele, knock-in band was gel-purified with Gel Extraction Kit (Qiagen). 200 ng of the gel-purified knock-in band was digested with SpeI enzyme (NEB) in a 10-μ l reaction. For the clones that had correct SpeI digestion pattern, we also did left and right junction PCRs. For the left junction PCR, we used the out-forward genotyping primer for each region and the “donor PCR Rev” primer. For the right junction PCR, we used “donor PCR For” primer and the out-reverse genotyping primer for each region. All the primer sequences are listed below. For all the gel images showing PCR results, we used either NEB 2-log or 1 kb ladder (NEB).

Sanger DNA sequencing was performed (Genewiz) to verify sequences of the left and right junctions as well as the integrity of the TetO repeats. For sequencing left and right junctions, “left junct Rev” and “right junct For”primers were used, respectively. For sequencing of the complete TetO repeat, “Seq tetO-mid For“, “Seq tetO-mid2 For” and “Seq tetO-mid Rev” primers were used as necessary.

Clone 10-22 had both alleles targeted and since the integration was through NHEJ, the alleles had differences in the sequence. To be able to sequence each allele separately, 10-22 knock-in band was amplified with a modified version of the forward and reverse genotyping primers to integrate 5’-BsmBI (giving AATT 5’ overhang) and 3“-BamHI sites, respectively. Knock-in band was digested with BsmBI and BamHI enzymes, followed by cloning into the pSP2 backbone from 48-mer donor plasmid through EcoRI and BamHI sites. Plasmids from single colonies were purified and sent for sequencing.

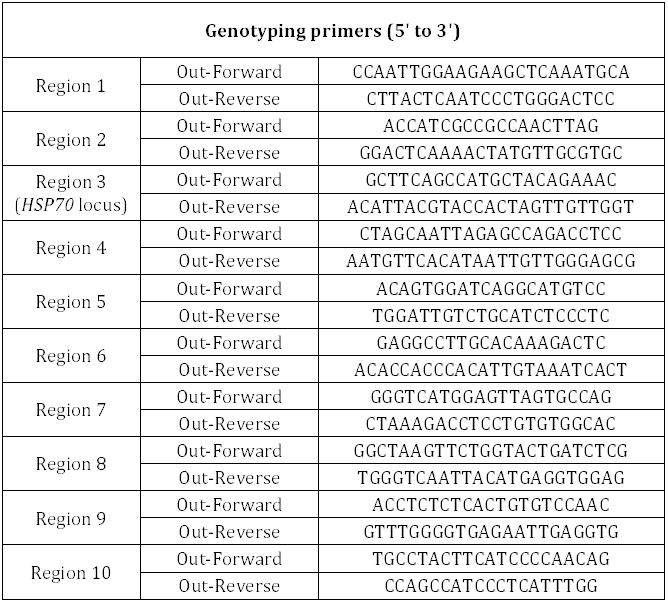
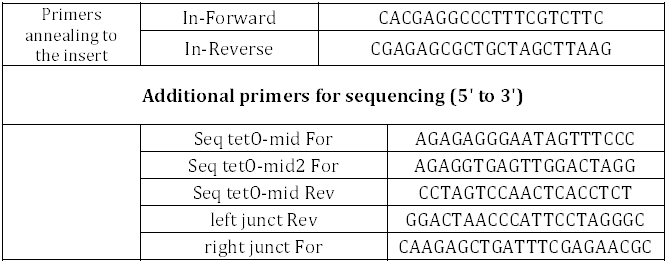

### Lentivirus Production and Transduction

For packaging, pCMV-VSV-G (Addgene plasmid # 8454) and pCMV-dR8.2 (Addgene plasmid # 8455) plasmids were used. One day before transfection, HEK293T cells were plated into 12-well plate and we aimed for ~ 50-60% confluency for the next day. For packaging, the following amounts of each plasmid were mixed: 0.5 μ g lenti F9-TetR-EGFP-ires-PuroR vector + 0.45 μ g pCMV-dR8.2 + 0.05 μ g pCMV-VSV-G. The plasmids were transfected using Fugene HD (Promega), following manufacturer’s protocol. After 24 hours, the media was changed into fresh DMEM + 10% FBS. ~ 60 hours after transfection, 1 ml viral supernatant was collected and filtered through a 0.45 μ m syringe filter. We avoided multiple freeze-thaw of the lentiviral supernatant.

For transduction, 1×10^5^ cells per well were plated into a 12-well plate with 8 μ g/ml polybrene (Santa Cruz Biotechnology, Dallas, TX). 100 μ l from the viral supernatant was added per well. Two days after transduction with the lenti F9-TetR-EGFP-ires-PuroR, puromycin selection (1 μ g/ml) was started. Cells were analyzed following 3 days of puromycin selection.

## RNAFISH

Before RNA FISH, clone 3-9-5 (expressing lenti F9-TetR-EGFP-ires-PuroR) was heat-shocked for 15 minutes at 42°C. 34 RNA FISH probes were designed (Biosearch Technologies, Novato, CA) and labeled with Cy5 at their 3’ ends. Cells grown on coverslips were fixed with 3.6% paraformaldehyde (Sigma-Aldrich) for 12 minutes at room temperature (RT). After three times of washing with calcium-magnesium free phosphate-buffered saline (CMF-PBS), cells were permeabilized in 0.5 % Triton X-100 for 15 minutes, rinsed three times with CMF-PBS, and then equilibrated in wash buffer (50% formamide and 2% SSC) for 30 minutes. Hybridization reaction was performed according to a previously published method (Raj et al., 2008). After 15 hours of hybridization, the cells were washed in wash buffer twice for 30 minutes, and mounted on a glass slide.

### Immunofluorescence Staining of the Knock-in Clones

HCT116 clone 3-9-5 and clone 10-21, which were infected with lenti F9-TetR-EGFP-ires-PuroR, were plated on collagen-coated coverslips (Neuvitro, Vancouver, WA) in a 24-well plate and grown to 80% confluency. Cells were fixed with 1.6% paraformaldehyde (PFA; Sigma-Aldrich) in phosphate-buffered saline (PBS) for 15 minutes at RT and redundant PFA was quenched with 0.125 M glycine for 5 minutes at RT. Cells were then permeabilized with 0.5% Triton in PBS for 15 minutes at RT, washed with 0.1% Triton (in PBS) for three times (each for 5 minutes) at RT, and blocked with 5% goat serum (G9023, Sigma-Aldrich) in 0.1% PBST (Phosphate-buffered saline with Tween 20, blocking buffer) for 1 hour at RT. Cells were then incubated overnight at 4°C with rabbit-anti-SON polyclonal (HPA023535, Sigma-Aldrich) and mouse-anti-LaminB1/B2 monoclonal Clone 2D8 (A gift from Robert Goldman) antibodies, each diluted 1:1000 in blocking buffer. Cells were washed with 0.1% PBST for three times (each for 5 minutes) at RT and then incubated for 5 hours at RT with Alexa Fluor^^®^^ 647-conjugated goat-anti-rabbit IgG and Alexa Fluor^®^ 594-conjugated goat-anti-mouse IgG (Jackson ImmunoResearch, West Grove, PA) antibodies, each diluted 1:200 in blocking buffer. Cells were then washed with 0.1% PBST for three times (each for 5 minutes) at RT. Coverslips were mounted in DAPI-containing, anti-fade mounting media.

### DNA-FISH Probe Labeling

Knock-in loci specific BACs (RP11-479I13 for region 3 and CTD-2643I7 for region 10) were used for preparing FISH probes. BACs (Invitrogen) were end labeled with terminal transferase with a modified protocol described elsewhere (Dernburg, 2011). Briefly, BAC DNA was digested into 100-150 bp fragments by incubating with a restriction enzyme cocktail including AluI, DpnI, HaeIII, MseI, MspI and RsaI (NEB), overnight at 37°C. DNA was purified by ethanol precipitation and denatured at 95°C for 5 minutes with snap chill. 2 µg of fragmented DNA was labeled with Terminal Deoxynucleotidyl Transferase (TdT, 20 U/μL) (Fermentas, Waltham, MA) in a 25 μl reaction with suitable oligonucleotides and the reaction was incubated at 37°C for 1 hour. RP11-479I13 was labeled with biotin by using 54 μM biotin-14-dATP (Thermo Fisher Scientific) and 108 μM dATP (NEB) in the labeling reaction. CTD-2643I7 was labeled with Dig by using 54 μM Dig-11-dUTP (Roche, Basel, Switzerland) and 108 μM dTTP (NEB) in the labeling reaction. Labeled probes were mixed with 10x (20 μg) human Cot-1 DNA (Invitrogen) in a final concentration of 2.5 M ammonium acetate (using glycogen (Roche) as a carrier), followed by precipitation with ethanol. Probes were then dissolved in hybridization buffer (10% dextran sulfate, 50% formamide, 2x SSC) with final concentration of 50 ngμl.

### 3D Immuno-FISH

3D immuno-FISH was conducted following a protocol modified from a previously published paper (Solovei and Cremer, 2010). Wt HCT116 cells were plated on collagen-coated coverslips in a 24-well plate and grown to 80% confluency. Cells were permeabilized in 0.1%Triton in PBS for 55 seconds at RT, fixed with 3.6% PFA in PBS for 10 minutes at RT, and quenched with 0.5 M glycine in PBS for 5 minutes at RT.

For immunostaining, cells were blocked with 5% goat serum (G9023, Sigma-Aldrich) for 1 hour at RT, followed by incubation with rabbit-anti-SON polyclonal (HPA023535, Sigma-Aldrich) and mouse-anti-LaminB1/B2 monoclonal Clone 2D8 (A gift from Robert Goldman) antibodies (each diluted 1:1000 in blocking buffer) overnight at 4°C. Cells were washed with 0.1% PBST for three times (each for 5 minutes) at RT, incubated for 5 hours at RT with FITC-conjugated goat-anti-rabbit IgG (Jackson ImmunoResearch; 1:200 diluted in blocking buffer) and AMCA-conjugated goat-anti-mouse IgG (Jackson ImmunoResearch; 1: 25 diluted in blocking buffer), and washed with 0.1% PBST for three times (each for 5 minutes) at RT. Cells were then fixed with 3% PFA in PBS for 10 minutes at RT, quenched with 0.5 M glycine in PBS for 5 minutes at RT, and washed with 0.1% PBST for 5 minutes at RT.

For *in situ* hybridization, coverslips were submerged into 20% glycerol (in PBS) for 60 minutes at RT, followed by 6 freeze-thaw cycles by liquid nitrogen. Coverslips were washed three times (each for 5 minutes) with 0.1% PBST, rinsed and incubated with 0.1N HCl/0.7% Triton/2x SSC for 15 minutes at RT, washed three times (each for 5 minutes) with 2x SSC, and incubated with 50% formamide/2XSSC for 30 minutes at RT. 50 ng/μl FISH probes were diluted by 1:3 with hybridization buffer, in a final volume of 6 μl, and then added onto coverslips and sealed onto glass slides with rubber cement. gDNA was denatured on a hot plate at 78°C for 3 minutes and then hybridized for 48 hours in a humid chamber at 37°C. Cells were washed with 2x SSC at 37°C for three times (each for 5 minutes), with 0.1x SSC at 60°C for three times (each for 5 minutes), and with 4x SSC at RT for 5 minutes. Cells were blocked in 4% BSA/4XSSC/0.1% Triton (blocking buffer) at RT for 1 hour and incubated overnight at 4°C with Streptavidin-Alexa Fluor ^^®^^ 594 (Invitrogen; 1:100 diluted in blocking buffer) and mouse-anti-DIG-Alexa Fluor^^®^^ 647 (Jackson ImmunoResearch; 1:200 diluted in blocking buffer). Cells were then washed in 4x SSC/0.1%Triton three times (each for 5 minutes) at RT and mounted in anti-fade mounting media.

### Image Acquisition and Analysis

For analysis of the immuno-FISH and immunofluorescence results, 3D optical-section images were acquired with 0.2 μ m z-steps using a DeltaVision OMX microscope system and DeltaVision SoftWorx software (GE Healthcare, Little Chalfont, UK) with a 100x, 1.4 NA objective lens and an Evolve 512 Delta EMCCD Camera with conventional light path. Images were deconvolved using an enhanced ratio, iterative constrained algorithm (Agard et al., 1989) and then registered based on Image Registration Target Slide (GE Healthcare) and TetraSpeck fluorescent beads (Molecular Probes, Eugene, OR). Image deconvolution and registration were done with the SoftWorx 6.5.2 software (GE Healthcare). The distributions were graphed in the program OriginPro (OriginLab, Northampton, MA).

Images from immunofluorescence staining of knock-in clones or immuno-FISH of wt HCT116 cells were analyzed to determine the distance of locus of interest from either nuclear speckles or lamina (Figure 6). The same analysis was also done for Movie 1 and 2 (Figure 7). The shortest distances were measured from the center of each EGFP or FISH spot to the edge of its nearest nuclear speckle or to the center of its nearest lamina region using the Straight Line tool in ImageJ (National Institutes of Health). All images in Figure 6 and 7 were prepared using ImageJ software.

To determine percentage of cells with paired loci and for analysis of the distance between the paired loci (Figure 5), a Structured illumination Superresolution system (Elyra SIM, Carl Zeiss, Obercohen, Germany) was used to analyze fixed cells. The system configuration was the same as described before (Sivaguru et al., 2016). Briefly, a 1.4 NA Plan Apochromat 63x objective with oil immersion was used together with a 488 nm excitation and the emission signals were picked up using an Andor Ixon EMCCD (EM gain setting of 50 and exposure time ~ 100 ms) camera between 500-550 nm bandpass filter. For each focal plane, five rotations and five phases were used and a z stack was made at 130 nm distance between slices and the final processed XY pixel size was 40 nm. The raw data was processed using the Zen software supplied by the manufacturer under Structured Illumination processing module with default (automatic processing) settings. The distance between paired spots was determined with the same program using the profile module and XY calipers between the two spots. Imaris program (Bitplane AG, Zurich, Switzerland) was used to generate the XY and YZ profiles to determine and display the single and double locus status across cell types. The distributions were graphed in the program OriginPro (OriginLab).

All other fixed samples were analyzed using Personal DeltaVision deconvolution microscope, which is equipped with Xenon lamp, 60X oil objective (NA 1.4) and CoolSNAP HQ slow-scan CCD camera (Roper Scientific, Vianen, Netherlands). After taking z-stack images, the image file was deconvoluted and projected into a 2D image using Softworx program (GE Healthcare).

For live cell imaging, we used the DeltaVision OMX microscope described above. Cells were grown in collagen-coated 35mm glass bottom dish (Mattek, Ashland, MA), and maintained at 37°C and 5% CO2 during imaging. Post-processing of the images was done as described above. Additionally, maximum projection and correction for the transformation of the nucleus (Thévenaz et al., 1998) were performed for the movie taken from the cells expressing nuclear speckles (Movie 1) by Softworx (GE Healthcare) and ImageJ (National Institutes of Health), respectively.

---

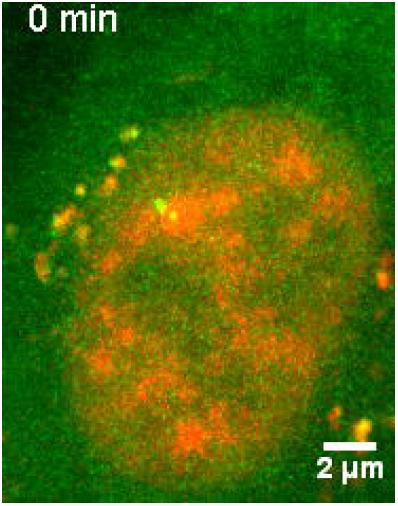

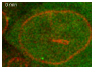

